# Pathfinder: A gamified measure to integrate general cognitive ability into the biological, medical, and behavioural sciences

**DOI:** 10.1101/2021.02.10.430571

**Authors:** Margherita Malanchini, Kaili Rimfeld, Agnieszka Gidziela, Rosa Cheesman, Andrea G. Allegrini, Nicholas Shakeshaft, Kerry Schofield, Amy Packer, Rachel Ogden, Andrew McMillan, Stuart J. Ritchie, Philip S. Dale, Thalia C. Eley, Sophie von Stumm, Robert Plomin

## Abstract

Genome-wide association (GWA) studies have uncovered DNA variants associated with individual differences in general cognitive ability (*g*), but these are far from capturing heritability estimates obtained from twin studies. A major barrier to finding more of this ‘missing heritability’ is assessment – the use of diverse measures across GWA studies as well as time and cost of assessment. In a series of four studies, we created a 15-minute (40-item), online, gamified measure of *g* that is highly reliable (alpha = .78; two-week test-retest reliability = .88), psychometrically valid and scalable; we called this new measure Pathfinder. In a fifth study, we administered this measure to 4,751 young adults from the Twins Early Development Study. This novel *g* measure, which also yields reliable verbal and nonverbal scores, correlated substantially with standard measures of g collected at previous ages (r ranging from .42 at age 7 to .57 at age 16). Pathfinder showed substantial twin heritability (.57, 95% CIs = .43, .68) and SNP heritability (.37, 95% CIs = .04, .70). A polygenic score computed from GWA studies of five cognitive and educational traits accounted for 12% of the variation in *g*, the strongest DNA-based prediction of *g* to date. Widespread use of this engaging new measure will advance research not only in genomics but throughout the biological, medical, and behavioural sciences.

## Introduction

Given its association with crucial life outcomes, it is essential to understand the genetic and environmental mechanisms that support the development of general cognitive ability (*g*). A major barrier in identifying the genetics of *g* is measurement heterogeneity. Traditional cognitive assessment is expensive and time-consuming and therefore unsuited to large biobanks; consequently, gene discovery studies have had to integrate data from multiple cohorts that differ widely in the quality of measurement of *g*. We present a brief, reliable, valid, and engaging new measure of *g, Pathfinder*, developed over four studies. In a fifth study we administered this measure to a large sample of young adult twins and assessed the psychometric and genetic properties of the measure.

General cognitive ability (*g*) is the best behavioural predictor of many educational, social and health outcomes (1). The symbol *g* was proposed more than a century ago to denote the substantial covariance among diverse tests of cognitive abilities. This underlying dimension runs through diverse cognitive abilities such as abstract reasoning, spatial ability and verbal ability and dominates the predictive validity of cognitive tests for educational, occupational, and life outcomes (2–4). In a meta-analysis of over 460 datasets, the average correlation among such diverse tests was about .30, and a general factor (first unrotated principal component) accounted for about 40% of the tests’ total variance (5).

Model-fitting analyses that simultaneously analyze the mountain of family, adoption, and twin data on g indicate that about half of the differences between individuals (i.e., variance) can be attributed to inherited DNA differences, a statistic know as *heritability* (6,7). Shared environmental influences that make family members similar to one another contribute 20% of the variance in parent-offspring studies, 25% in sibling studies and 35% in twin studies (6). However, one of the most interesting and perhaps counterintuitive findings about *g* is the developmental change in these estimates. Heritability increases from 45% in childhood to 55% in adolescence to 65% in adulthood, while shared environmental influence decreases from 30% to 15% in twin studies (7,8) and is even less in adoption studies (9).

Multivariate genetic analysis, which examine associations between multiple traits, shows that genetic overlap among cognitive tests is much greater than their phenotypic overlap. The average genetic correlation among diverse cognitive tests is about .80, indicating that many of the same genes affect different cognitive abilities (10–12). Recent evidence applying genomic methods has shown that this genetic covariance is largely reflected in the *g* factor (12).

Progress in identifying some of the many DNA differences that account for the heritability of *g* would result in advances not only in genomics, but across the psychological, biological and medical sciences (13). This is because *g* pervades virtually all aspects of life, including education (14), job satisfaction and earnings (15,16) and health and longevity (17–20). A substantial portion of the observed associations between *g* and education, wealth and health is rooted in genetic variation (21,22). For example, substantial genetic correlations have been observed between *g* and educational attainment (*r* = 0.73), longevity (*r* = 0.43) and age of first birth (*r* = 0.46; (23)). This widespread pleiotropy (i.e. the same genetic variants contributing to two or more traits) suggests that *g* can be a useful translational target for any area of research in the life sciences –biology, brain as well as behaviour (24).

Given the genetic overlap observed between *g* and physical and mental health (25), advances in uncovering the DNA variants associated with individual differences in *g* are likely to enhance our understanding of the genetics of health, illness and psychiatric disorders. This becomes particularly meaningful when considering the major challenges related to gene discovery in specific areas of the psychological and medical sciences, most prominently psychiatric disorders (26). With the notable exception of schizophrenia, for which a polygenic score constructed from the latest GWA study (27) was found to account for 7.7% of the variance in liability in independent samples, genomic prediction of psychiatric traits and disorders has been considerably less successful (28,29) than for *g* (25,30). Leveraging on pleiotropy, progress in uncovering the genetics of *g* might therefore exert important spillover effects for our understanding of the genetics of physical and mental health.

We now know that the biggest effects of specific DNA variants associated with most complex traits, including *g*, account for less than 0.1% of the variance (31). Genome-wide association (GWA) studies that attempt to identify these DNA associations need very large samples to reliably detect the tiny effects; however, testing large samples on *g* is challenging. As a result, it has been necessary to meta-analyze GWA results across studies that have used different methods and measures to assess *g*.

The largest meta-analytic GWA study of *g* included a total of 270,000 individuals from 14 cohorts, all of which used different measures of *g* (23). Despite the heterogeneity of measures, this GWA study was able to identify 242 independent loci significantly associated with variation in *g*. A polygenic score derived from this GWA meta-analysis predicted 7% of the variance in *g* at age 16 in the sample used in the present study (30). A polygenic score for *g* is a genetic index of *g* for each individual that represents the sum across the genome of thousands of DNA differences associated with *g* weighted by the effect size of each DNA variant’s association with *g* in GWA studies. Adding a polygenic score derived from a GWA meta-analysis for years of schooling (32) to the polygenic score for *g* boosts the prediction of *g* to 10% at age 16 (30).

Nonetheless, 10% is a long way from the heritability estimate of 50% from twin studies. This gap is known as ‘missing heritability’, which is a key genetic issue for all complex traits in the life sciences (33). Increasing GWA sample size and employing whole-genome sequencing approaches that can capture rare variants are among the approaches in use to narrow the missing heritability gap (34). Better measurement of the phenotype can also help. Differences between the psychometric quality of measures have been shown to reduce the statistical power to detect genetic associations, the effect sizes of the detected associations, and the predictive power and specificity of the polygenic scores that derive from GWA studies (36–38). For example, a simulation study showed that with heterogeneity of 50%, the sample size needed to achieve the same statistical power obtained from homogeneous samples increased by approximately three times (36). Extant GWA studies of *g* differ widely in the quality of measurement, from individually administered full-scale IQ tests to scores on a college entrance exam or a single reading test or six items on a digit-span test (Savage et al., 2018). Rather than combining small heterogeneous GWA studies with diverse measures of *g*, a better strategy is to incorporate the same high-quality measure of *g* in large biobanks that already have genotype data on their participants. Cognitive testing has not been conducted in most biobanks because traditional in-person testing is expensive and time-consuming.

This issue of heterogeneity of measurement in GWA studies motivated us to create a brief, reliable and valid online measure of *g* that could be offered to participants in extant biobanks. In addition to the criterion of brevity (15-minute) and ease of access and use, we set out to develop a *g* measure characterized by an additional important feature: gamification. Evidence points to the positive impact of gamification on participants’ engagement and motivation (39,40), which boosts the value of on-line gamified tests, for two reasons. First, increasing engagement and motivation is likely to reduce distractions and drift in attention, leading to more reliable estimates of performance, especially in online testing conditions outside the controlled environment of the laboratory. Second, participants’ satisfaction increases participation and retention rates, which is especially important for large cohort studies (41).

Gamification sets our measure apart from the few other existing online batteries that are capable of reliably assessing *g*. The two most prominent examples are the battery of cognitive tests that has been developed for and administered to UK Biobank participants (42) and the Great British Intelligence Test (43), a citizen science project launched in late December 2019 by BBC2 Horizon. The Great British Intelligence Test includes a selection of 9 cognitive tests from a broader library of 12 tests available via the *Cognitron* repository, which takes 20-30 minutes to complete. The cognitive tests administered to UK Biobank participants, which take on average 21 minutes to complete, assess five abilities: visual memory, processing speed, numeric working memory, prospective memory, and verbal and numerical reasoning. Recent analyses found the tests to have moderate concurrent validity, with a mean correlation between the shortened version and a validated reference test of 0.53, but ranging widely from 0.22 to 0.83, and moderate short-term stability, with a mean four-week test-retest correlation of 0.55, ranging between 0.40 and 0.89 for individual tests (44). In addition, although the five tests yielded a measure of *g* that correlated 0.83 with a measure of *g* constructed from their corresponding standardized reference tests, the estimate of *g* provided by the battery appears to reflect the fluid, largely not dependent on prior learning, aspects of intelligence more strongly than the crystallized, academic forms of cognitive function, such as vocabulary and verbal knowledge (12).

Our *g* battery overcomes these limitations by providing a highly reliable, balanced assessment of *g*, constructed from measures of verbal and nonverbal abilities. Importantly, and different from all existing measures, our measure is gamified and engaging, accessible by all researchers through our open science research framework, and easy to integrate within existing data collection platforms. It is also at least five minutes shorter than existing measures, which is particularly meaningful when considering data collection in large cohorts.

The current paper describes our work developing and validating this new, brief, easy-to-administer, gamified measure of *g* in a series of four studies (see **Figure 1**). In a fifth study, we administered this measure to 4,751 young adults from the Twins Early Development Study (TEDS; Rimfeld et al., 2019) and assessed the psychometric and genetic properties of the measure. Our analyses were preregistered in line with the Open Access Framework (https://osf.io/pc9yh/), and included the following three core hypotheses:

**Figure 1.**
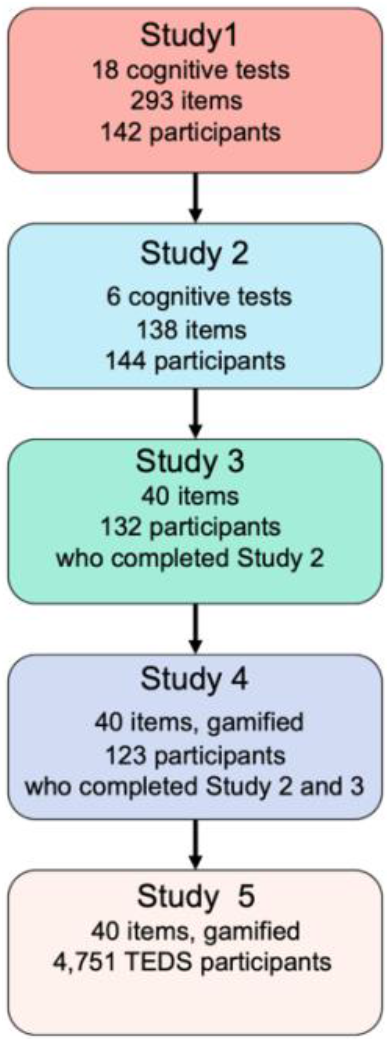
Flowchart depicting the roadmap to the development and testing of Pathfinder over our five studies.

First, we hypothesized that our 15-minute online measure of *g* would:

a. Capture more than 40% of the variance of diverse tests of verbal and nonverbal abilities in a first principal component.
b. Yield test-retest reliability greater than 0.80. Second, we predicted that, using the classical twin design, our measure of *g* would:
c. Yield heritability estimates greater than 50%.
d. Yield estimates of shared environmental influence less than 20%.

Third, we predicted multivariate approaches to calculating polygenic scores would predict over 10% of the variance in individual differences in *g* in our sample of young adults.

## Materials and Methods

### Study 1

#### Participants

Participants (N = 142) were recruited from the Twins Early Development Study (TEDS) sample(46). Specifically, for this first study we invited a group of TEDS twins whose co-twin was no longer actively participating in the TEDS longitudinal data collection. Sixteen out of the 142 participants who agreed to take part in the study did not complete the full battery, which resulted in N = 126 participants with complete data. Participants’ ages ranged between 21.60 and 22.30 (M = 21.98, SD = .19). The sample included more females (*N* = 97) than males (*N* = 45). Participants varied in their education level (58% had completed A-level exams).

#### Measures

##### Cognitive battery

Participants were administered a battery of 18 well-established cognitive tests covering four core domains of cognitive performance, including a total of 293 items: Nonverbal reasoning (6 tests for a total of 75 items), Verbal reasoning (4 tests for a total of 98 items), Spatial ability (3 tests for a total of 45 items) and Memory (5 tests for a total of 75 items, 2 tests assessed long-term memory and 3 tests short-term memory). A full list of tests is reported in **Supplementary Table 1** and examples for each test can be found at the following link: https://www.youtube.com/watch?v=TA38bsgp7Lg&ab_channel=TEDSProject. The 18 tests were selected after a careful literature review and were chosen with three core features in mind: (1) each test had to demonstrate high validity and reliability; (2) altogether, tests had to tap a wide array of cognitive domains, from verbal and nonverbal reasoning to memory; and (3) they had to be tests that were either developed or adapted for online administration, or tests that could easily be adapted by our team for online administration.

The final battery was administered online using forepsyte.com, an online data collection platform. Tests were presented in a fixed order and the order of presentation is reported in **Supplementary Table 1**. The median time participants took to complete the battery was 68 minutes.

### Study 2

#### Participants

Participants (*N* = 144) were recruited using Prolific.co (www.prolific.co), an online research recruitment platform. Of the total sample, 30% (*n* = 43) were males, 68% (*n* = 98) females, and 2% (*n* = 3) did not specify their gender. Participants’ ages ranged from 18 to 49 years (M = 30.99, *SD* = 8.67). Recruitment was based on four selection criteria: 1) age between 18 and 50 years; 2) English as first language; 3) UK nationality; and 4) education level which was selected in two groups, one of which had completed tertiary education and the other not (this resulted in 40.9% of the total sample who had completed tertiary education and 59.1% not, which is representative of educational levels in the UK population; seehttps://www.oecd.org/unitedkingdom/United%20Kingdom-EAG2014-Country-Note.pdf). **Supplementary Table 2** presents a breakdown of the participants’ education level and ethnicity.

#### Measures

##### Cognitive battery

The cognitive battery included 138 items (78 verbal and 60 non-verbal) from seven well-established cognitive tests, which were selected from a larger battery based on the results of Study 1. The three tests assessing verbal ability were: (1) the Mill Hill Vocabulary Scale (47), (2) a Missing Letter Test and (3) a Verbal Analogies Test. The <u>Mill Hill Vocabulary Scale</u> consists of items assessing individuals’ ability to select semantically related words. For each item a target word is displayed, and participants are asked to select the word that is closest in meaning from six response options. In the <u>Missing Letter Test</u>, participants were exposed to pairs or strings of words, each with a blank space indicating a missing letter. Participants were asked to identify the missing letter that would meaningfully complete all the words presented on the screen simultaneously and select the letter from a displayed alphabet. An example of items is *ban(?) (?)ave – fla(?) (?)ain* and the missing letter in this instance is “g”. In the <u>Verbal Analogies Test</u>, participants were presented with verbal analogies, having either one or two missing words. An example of a one-word problem is: “Sadness is to happiness as defeat is to *x*”. Participants could solve *x* by choosing between four options: Joy, Victory, Victor, Tears. An example of a two-word problem is: “Robin is to *x* as Spider is to *y*”. Participants could choose between four options to solve *x* (Batman, Bird, Christmas, Tree) and four options to solve *y* (Spiderman, Easter, Arachnid, Insect). Participants were asked to select the word(s) that would correctly and meaningfully complete the missing part of the sentence. For items containing one missing word, participants selected their answer from a choice of four or five. For items with two missing words a choice of four was presented for every word missing.

The four tests assessing nonverbal ability were: (1) the Raven’s Standard Progressive Matrices (48), and three Visual Puzzles tests: (2) Non-verbal Analogies, (3) Non-verbal groupings and (4) Nonverbal Logical Sequences. The Raven’s progressive matrices test measures non-verbal abstract reasoning. Participants are presented with a series of incomplete matrices and are asked to select the missing part from a choice of eight. In the non-verbal analogies test, participants are presented with a series of images that contain a logical statement phrased as “*x* is to *y* as *z* is to_“, where *x/y/z* are replaced by images. Participants are asked to select the correct missing image to complete this statement. In the non-verbal groups test, participants are presented with the image of a group of shapes and are asked to identify which other shape, out of five options, belongs to the group. In the non-verbal sequences test, participants are presented with items containing a sequence of images, in which one is removed and replaced by a question mark and they are asked to select the image that completes the sequence from five options.

The seven tests (three verbal and four nonverbal) were presented to participants in a randomized order. Within each test, items were presented in fixed order, starting from easier items (determined from the results of study 1) and moving on to progressively more difficult ones. Each item was presented for a maximum of 60 seconds.

### Study 3

#### Participants

About two weeks (mean = 13.00 days) after the completion of Study 2, participants were invited back to participate in Study 3. Of those invited back, 91.7% completed Study 3 (*N* = 132). Out of the total sample for Study 3, 30.3% (*n* = 40) were males, 67.4% (*n* = 89) were females, and 2.3% (*n* = 3) did not specify their gender; the mean age was 31.3 years (*SD* = 8.7), and age ranged between 18 and 49 years. **Supplementary Table 2** presents a breakdown of the participants’ education levels and ethnicities.

#### Measures

##### Cognitive battery

The cognitive battery included the 40 items selected based on the results of Study 2. These 40 items covered five tests: 3 capturing verbal ability (Vocabulary, Verbal analogies and Missing letter) and 2 nonverbal ability (Matrix reasoning and Visual puzzles, the latter being a composite of the best-performing items from each of the three visual puzzles tests administered in Study 2). The order of presentation of these tests was randomized to account for the potential effects of test-taking fatigue on cognitive performance. Within each test, items were presented in order of difficulty, based on accuracy results from Study 2 (**see Supplementary Table 3**). Each item was presented for between 20 to 40 seconds, the time limit decisions were made based on the means and standard deviations for response time obtained from Study 2 (**see Supplementary Table 3**). During this phase we also added four quality control (QC) items. These were presented in the same form as test items, but they were extremely easy to solve; their aim was to help us identifying ‘clickers’, i.e., participants who were just clicking through the test and providing random responses. Control items did not contribute to either the tests or total score. A fifth standard quality control question ‘*This is a quality control question, please select option B*’ was also added. QC items were presented half-way through each test, except for the standard quality control question that was presented between two tests in randomized order. Response accuracy for each QC item is presented in **Supplementary Table 4**.

### Study 4

#### Participants

Approximately one month after Study 3 (mean = 29, range = 23 to 35 days), participants who completed both Study 2 and Study 3 were invited back to complete Study 4. Of those invited back, 85.4% completed Study 4 (*N* = 123). Out of the total sample for Study 4, 30.1% (*n* = 37) were males, 68.3.% (*n* = 84) were females, and 1.6% (*n* = 2) did not specify their gender; the mean age was 31.82 years (*SD* = 8.61), and age ranged between 18 to 50 years. **Supplementary Table 2** presents a breakdown of the participants’ education levels and ethnicities.

#### Measures

##### Gamified cognitive battery

In Study 4 we administered the same battery of 40 items included in Study 3, but this time the items were embedded in a gamified storyline, the *Pathfinder*, which took participants through five ‘journeys’. A detailed description of each journey can be found in the TEDS data dictionary at the following link: http://www.teds.ac.uk/datadictionary/studies/webtests/21yr_ggame_description.htm. **Figure 2** provides a visual summary of the graphics of how items were presented in the gamified test and **Figure 2F** provides an example of the feedback that participants were given at the end of the gamified test.

**Figure 2.**
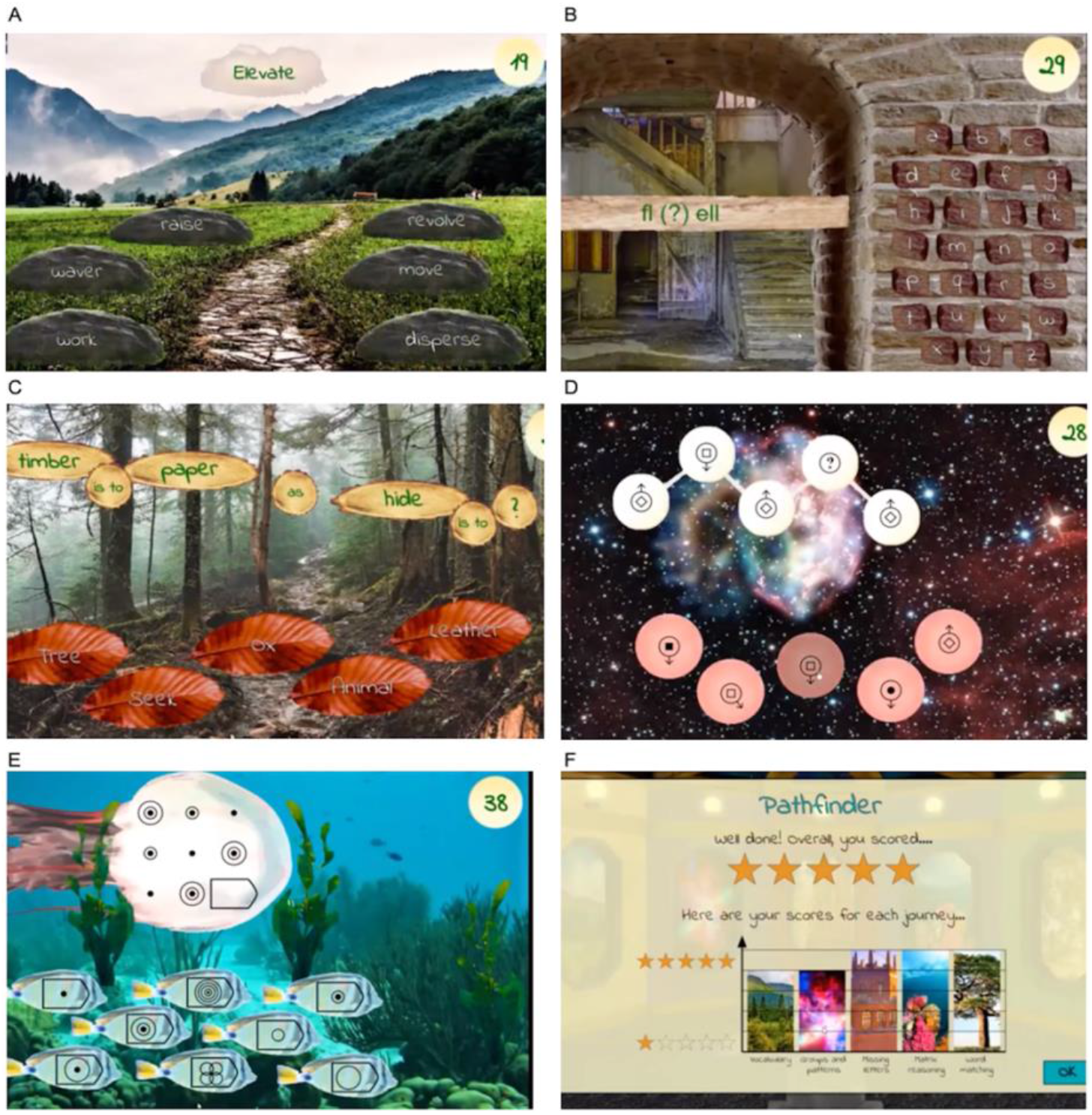
Screenshots of each of the five ‘journeys’ included in the *Pathfinder* gamified test (panels A-E) and a visual representation of the final feedback page (panel F). Panel A depicts the “Mountain” journey (Vocabulary test); panel B the “Tower” journey (Missing letter test); panel C the “Woodland” journey (Verbal analogies test); panel D the “Space” journey (Visual puzzles); and panel E the “Ocean” journey (Matrix reasoning test).

### Study 5

#### Participants

In study 5, *Pathfinder* was administered to an initial sample of 4,751 twins (1,491 twin pairs and 1,769 individual twins) from the Twins Early Development Study (TEDS) (45). All families with twins born in England and Wales between 1994 and 1996, identified through birth records, were invited to take part in TEDS. Over 15,000 families took part in the first data collection wave and over 10,000 families are still actively participating in TEDS 25 years on. TEDS is an ongoing project and TEDS twins have contributed data longitudinally from birth to the present day. The last major wave of assessment was conducted in 2018 when the twins were 21-23 years old. TEDS remains reasonably representative of the UK population in terms of ethnicity and socioeconomic status (SES; see (46) for a detailed description). Data from twins known to suffer from a severe medical condition including autism, cerebral palsy, chromosomal or single-gene disorders and organic brain problems, were excluded from the current analyses, together with twins whose sex and/or zygosity was unknown (N = 137 participants excluded). In addition, ‘clickers’ were identified from a combination of the incorrect responses in QC items, rapid responding (based on the mean item response time), low sub-test score and uniform responding (i.e. a pattern of clicking on the same response over a series of items). This resulted in the exclusion of data from 69 additional participants. The final sample consisted of 4,545 participants (1,416 twin pairs –639 monozygotic and 777 dizygotic pairs, and 1,713 unpaired twins). The sample mean age was 24.81 (SD = 0.85), ranging between 23.29 and 26.41. Genotyped DNA data was available for a subsample of 1,365 unrelated individuals. Genotypes underwent phasing using EAGLE2 and imputation into Haplotype Reference Consortium (release 1.1.), employing the Positional Burrows-Wheeler Transform method via the Sanger Imputation Service (see (49) for additional information). TEDS data collections have been approved by the King’s College London ethics committee.

#### Measures

##### Pathfinder

In Study 5 we administered the same tests administered in Study 4. The 15-minute (median time taken to complete the battery = 15.95 minutes), gamified *Pathfinder g* measure included two core components assessing verbal and nonverbal cognitive ability. The verbal ability block included 20 items from 3 tests: vocabulary, verbal analogies and missing letter. The nonverbal ability block included 20 items from 2 tests: matrix reasoning and visual puzzles (which grouped items from three tests: non-verbal analogies, non-verbal groupings and nonverbal logical sequences). The items were embedded in a gamified storyline as participants solved puzzles while moving through different journeys (represented as background images, which changed after every 1-3 items): The “Mountain” journey (Vocabulary test) included 8 items, the “Tower” journey (Missing letter) included 6 items, the “Woodland” journey (Verbal analogies) included 6 items, the “Space” journey (Visual puzzles) included 9 items, and the “Ocean” journey (Matrix reasoning) included 11 items (see **Figure 2**). The test included the same 5 QC items described in Study 3 and included in Study 3 and 4. ***Screen size***. Participants could complete *Pathfinder* using a variety of devices, including laptops, tablets and mobile phones. To account for the potentially confounding effects of screen size we created a categorical variable reflecting three screen size categories in order to statistically control for the effects of screen size. These categories were “small screen (< 768 pixels)”, “medium screen (768-1199 pixels)” and “large screen (>=1200 pixels)”.

##### Cognitive ability at earlier ages

TEDS includes measures of cognitive ability collected at multiple waves from childhood to late adolescence.

At **age 7** cognitive ability was measured using four cognitive tests that were administered over the telephone by trained research assistants. Two tests assessed verbal cognitive ability: a 13-item Similarity test and 18-item Vocabulary test, both derived from the Wechsler Intelligence Scale for Children (WISC; (50)). Nonverbal cognitive ability was measured using two tests: a 9-item Conceptual Groupings Test (51), and a 21-item WISC Picture Completion Test (50). Verbal and nonverbal ability composites were created taking the mean of the standardized test scores within each domain. A *g* composite was derived taking the mean of the two standardized verbal and two standardized nonverbal test scores.

At **age 9** cognitive ability was measured using four cognitive tests that were administered as booklets sent to TEDS families by post. Verbal ability was measured using the first 20 items from WISC-III-PI Words test (52) and the first 18 items from WISC-III-PI General Knowledge test (52). Nonverbal ability was assessed using the Shapes test (CAT3 Figure Classification; (53) and the Puzzle test (CAT3 Figure Analogies; Smith et. al., 2001). Verbal and nonverbal ability composites were created taking the mean of the standardized test scores within each domain. A *g* composite was derived taking the mean of the two standardized verbal and two standardized nonverbal test scores.

At **age 12**, cognitive ability was measured using four cognitive tests that were administered online. Verbal ability was measured using the full versions of the verbal ability tests administered at age 9: the full 30 items from WISC-III-PI Words test (52) and 30 items from WISC-III-PI General Knowledge test (52). Nonverbal ability was measured with the 24-item Pattern test (derived from the Raven’s Standard Progressive Matrices; (54) and the 30-item Picture Completion test (WISC-III-UK) (50). Verbal and nonverbal ability composites were created taking the mean of the standardized test scores within each domain. A *g* composite was derived taking the mean of the two standardized verbal and two standardized nonverbal test scores.

At **age 16** cognitive ability was measured using a composite of one verbal and one nonverbal test administered online. Verbal ability was assessed using an adaptation of the Mill Hill Vocabulary test (47), Nonverbal ability was measured using an adapted version of the Raven’s Standard Progressive Matrices test (47). A *g* composite was derived taking the mean of the two standardized tests.

##### Academic achievement at earlier ages

Measures of academic achievement have been obtained in TEDS throughout compulsory education.

At **age 7** academic achievement was measured with standardized teacher reports and consisted of standardised mean scores of students’ achievements in English and mathematics, in line with the National Curriculum Levels. Performance in English was assessed in four key domains: speaking, listening, reading and writing abilities; performance in maths was assessed in three key domains: applying mathematics, as well as knowledge about numbers, shapes, space and measures.

At **age 9**, academic achievement was again assessed using teacher reports. The domains assessed were the same for English and mathematics (although on age-appropriate content). In addition, performance in science was assessed considering two key domains: scientific enquiry and knowledge and understanding of life processes, living things and physical processes.

At **age 12**, academic achievement was assessed in the same way as at age 9, with the exception of mathematics, which was added a fourth domain: data handling, and science, which added a third domain: materials and their properties; these additions were in line with the changes made to the to the National Curriculum teacher ratings.

At **age 16**, academic achievement was measured using the General Certificate of Secondary Education (GCSE) exam scores. The GCSEs are the UK nationwide examination usually taken by 16-year-olds at the end of compulsory secondary education (55). Twins’ GCSE scores were obtained via mailing examination results forms to the families shortly after completion of the GCSE exams by the twins. For the GCSE, students could choose from a wide range of subjects. In the current analyses the mean score of the compulsory GCSE subjects English Language and/or English Literature, mathematics and a science composite (a mean score of any of the scientific subjects taken, including physics, chemistry and biology).

At **age 18**, academic achievement was measured based on the A-Level (Advanced Level) grade. The A-Level is a subject-based qualification conferred as part of the General Certificate of Education, as well as a school leaving qualification. A Levels have no specific subject requirements. We used standardized mean grade from all of the A-levels taken. Sample size was limited to those twins who who continued with academic education beyond GCSE level, typically in preparation for university, thus reducing range as well.

##### Family socioeconomic status (SES)

At **first contact**, parents of TEDS twins received a questionnaire by post, and were asked to provide information about their educational qualifications and employment and mothers’ age at first birth. SES was created by taking the mean of these three variables standardized. The same measures, except for mother’s age at first birth, were used to assess SES at **age 7**. At **age 16**, the SES was assessed based on a web questionnaire, and comprised a standardized mean score obtained from 5 items: household income, mother’s and father’s highest qualifications, mother’s and father’s employment status.

### Analyses

#### Phenotypic analyses

Phenotypic analyses were conducted in R version 4.0 (R Core Team, 2020) and Mplus version 8 (56). The variables were adjusted for the effects of sex, age (and screen size for the Pathfinder measures) using linear regression. Sex and age-controlled data were used in all downstream analyses. Because of the normal distribution of the *Pathfinder* measures no transformations were applied.

We conducted univariate <u>analysis of variance (ANOVAs)</u> to explore phenotypic sex differences and <u>Pearson’s correlations</u> to examine phenotypic associations between measures. We conducted <u>Principal Component Analysis (PCA)</u> and <u>Confirmatory Factor Analysis (CFA)</u> to examine the factor structure of the Pathfinder measures.

We applied <u>Item Response Theory (IRT)</u> modelling to reduce the cognitive battery, selecting items based on their psychometric properties. IRT refers to a set of mathematical models that describe the relationship between an individual’s response to test items and their level of the latent variable being measured by the scale – in this case, *g*. IRT allows estimation of item information, difficulty, and discrimination parameters (57). An item’s information properties are reflected in its item information curve, and its difficulty and discrimination properties are reflected in its item characteristic curve. *Item information* reflects the reliability of an item at a particular level of latent ability. The flatter the item information curve, the less reliable the item. An information curve positioned further along the x-axis suggests that an item is informative at the upper end of latent ability. *Item difficulty* is the level of latent ability at which the probability of correct response is 50%. The more difficult the question, the further the item characteristic curve will be to the right (more latent ability is needed to get it correct). *Item discrimination* indicates how much an item is influenced by the latent trait and is thus similar to a factor loading. High discriminative ability is indicated by a steep item characteristic curve. An item discriminates well at a particular level of *g* if a small change in ability results in a large increase in the probability of correct response. We fitted a binary 2-PL Model in the MPLUS software including all 138 verbal and non-verbal items. This model uses maximum likelihood and estimates item difficulty and discrimination (whereas the 1 PL model assumes items are equally discriminative). The 2 PL model provided a better fit for the data (Akaike Information Criterion (AIC) = 18327.596, Bayesian Information Criterion (BIC) = 19158.532, sample-size adjusted BIC= 18285.045) than a three-item parameter (3 PL) model (AIC = 18496.790, BIC = 19743.193, sample-size adjusted BIC= 18432.962), as indicated by the lower AIC, BIC and sample-size adjusted BIC indices obtained for the 2 PL IRT model, and a 1PL model which failed to converge.

#### Genetic and genomic analyses

##### The twin method

We applied the <u>univariate twin method</u> to partition the variance in each phenotype into genetic, shared and unique environmental influences. The twin method capitalizes on the genetic relatedness between monozygotic twins (MZ), who share 100% of their genetic makeup, and dizygotic twins (DZ), who share on average 50% of the genes that differ between individuals. The method is further grounded in the assumption that both types of twins who are raised in the same family share their rearing environments to approximately the same extent (58). By comparing how similar MZ and DZ twins are for a trait (intraclass correlations), it is possible to estimate the relative contribution of genetic and environmental factors to individual variation. Heritability (h^2^), the amount of variance in a trait that can be attributed to genetic variance (A), can be roughly estimated as double the difference between the MZ and DZ twin intraclass correlations (59). The ACE model further partitions the variance into shared environment (C), which describes the extent to which twins raised in the same family resemble each other beyond their shared genetic variance, and non-shared environment (E), which describes environmental variance that does not contribute to similarities between twin pairs (and also includes measurement error). Structural equation modelling provides more formal estimates of A, C, and E and calculates confidence intervals for all estimates. We performed twin analyses using OpenMx 2.0 for R (60) and Mplus version 8 (56).

Model fit was measured using the difference between the likelihood (−2LL) of the assumed model (with fewer parameters) and the likelihood of the saturated model, which provides a baseline summary of the data prior to decomposition into variance components (61). Difference in -2LL is distributed as chi-square (χ^2^) with χ^2^degrees of freedom (df) representing the difference in number of parameters between the baseline and more restrictive models. χ^2^ and df are used to create a p value for model fit comparisons, with a non-significant p-value indicating that the more restrictive model does not fit the data significantly worse than the saturated model (61).

The twin method was then extended to the exploration of the covariance between pairs of traits (<u>bivariate twin models</u>), by modelling cross-twin cross-trait covariances. Cross-twin cross-trait covariances describe the association between two variables, with twin 1’s score on variable 1 correlated with twin 2’s score on variable 2, which are calculated separately for MZ and DZ twins. We employed the bivariate twin models to explore genetic and environmental overlap between the *Pathfinder* composites and educationally relevant traits over development, using OpenMx 2.0 for R.

##### SNP heritability (SNP h^2^)

SNP heritability was estimated using the Genome-wide complex trait analysis (GCTA) software that employs a genome-based restricted maximum likelihood method (GREML). GREML estimates the proportion of the variance in a trait that is captured by all genotyped single nucleotide polymorphisms (SNPs) in samples of unrelated individuals (62). GREML uses individual-level genotypic data to estimate narrow-sense SNP h^2^, the proportion of phenotypic variation explained by the additive effects of genetic variants measured using a genotype array and subsequent imputation (62). Cryptic relatedness was controlled for by setting the relatedness threshold to .05, which resulted in removing pairs of individuals who are genetically as similar as 4th-degree relatives (63). The grm-adj 0 option was used to control for incomplete tagging of causal variants. Due to the fact that causal regions are likely to show lower MAF (minor allele frequency) compared to the genotyped set of genetic variants, weak LD (linkage disequilibrium) estimates may result. Incomplete tagging of causal loci may therefore be mitigated by assuming similar allele frequencies of causal loci and genotyped SNPs (63).

##### Genome-wide polygenic scores (GPS)

We constructed GPS using LD-pred (64) with an infinitesimal prior, which corrects for local linkage disequilibrium (LD), correlations between SNPs. We used the 1000 genomes phase 1 sample as a reference for the LD structure (see (65) for a detailed description of LD-pred analytic strategies used in calculating GPS in the TEDS sample). Three univariate polygenic scores were calculated from GWA summary statistics of intelligence (IQ3; N= 266,453 (23)), years of education (EA3; excluding 23andMe; N= 766,345 (66)) and childhood IQ (N= 12,441;(67)). Because the original IQ3 GWA meta-analysis included the TEDS sample, we used summary statistics that excluded TEDS to avoid bias due to sample overlap. The EA3 summary statistics employed here do not include 23andMe data (∼300k individuals) due to their data availability policy.

In addition to examining the predictions from individual GPS, we investigated the extent to which multivariate approaches boost the GPS prediction of g, verbal and nonverbal ability. Following the pipeline developed by Allegrini et al. (2019), multivariate polygenic scores were constructed using MTAG (68) and Genomic SEM (69), and combined the IQ3 and EA3 GPS with summary statistics of three additional educationally relevant traits: household income (N= 96,900; (70)), age at completion of full-time education (N= 226,899; (69)) and time spent using computer (N= 261,987; (71)).

Linear regression analyses were performed in R to investigate the association between the GPS and *Pathfinder* composites (R Core Team, 2017). We report results for the GPS constructed assuming a fraction of casual markers of 1 (assuming that all markers have non-zero effects). GPS results for other fractions (p-value thresholds) are included in the Supplementary Material. Phenotypic data, polygenic scores and covariates were standardized prior to the regression analysis to achieve the z-distribution and obtain R^2^ estimates in units of standard deviation. Variance explained by the GPS was determined as the difference between variance explained by the full model (including both GPS and covariates as predictors) and the null model (including the covariates alone). Each linear regression analysis included the following covariates: batch, chip and 10 principal components of population structure. All analyses were performed on samples of unrelated individuals.

## Results

Over four studies we adopted multiple psychometric approaches to develop the shortest possible, yet highly valid and reliable, measure of general cognitive ability (*g*).

### Study 1: Identifying the most informative verbal and nonverbal cognitive tests: Principal component analysis

In study 1 we administered a battery of 18 widely used cognitive tests, which we identified through an in-depth review of the literature. The sample and procedures are detailed in the Methods section. The final battery included 293 items that spanned four key areas of cognitive performance: nonverbal reasoning (75 items), verbal reasoning (98 items), spatial ability (45 items) and memory (75 items). **Supplementary Table 1** presents a full list of tests, which are described in greater detail in the Methods section, a demonstration of each test is provided at the following link: https://www.youtube.com/watch?v=TA38bsgp7Lg&ab_channel=TEDSProject. We conducted Principal Component Analysis (PCA) of these 18 tests to reduce the number of tests and select those that best represent verbal and nonverbal cognitive ability, the two core subdomains of cognitive skills which also reflect the key distinction between verbal and performance IQ.

We ran two separate PCAs, one for the 12 nonverbal measures and a second for the six verbal measures. The first PCA (**Supplementary Table 5a**) identified four tests that most reliably captured nonverbal reasoning, indexed by the highest loadings onto the first principal component (PC) of nonverbal ability. These nonverbal tests assessed Matrix reasoning (Raven’s progressive matrices), and Visual puzzles (Groups, Sequences and Nonverbal analogies). The second PCA (**Supplementary Table 5b**) indicated three tests that captured the majority of the variance in verbal ability: Similarities (Verbal analogies), Vocabulary (Mill Hill vocabulary test) and Information (Missing letter test). A first principal component including these seven tests accounted for 60% of the total variance. A composite *g* score (the scores summed) created from these seven tests, including a total of 138 items (average correlation across all individual items = .10), was strongly correlated (*r* = .85, p < .001, N = 126) with a g composite constructed from the entire battery (293 items). Cronbach’s alpha for each of the seven tests is reported in **Supplementary Table 5c;** the average alpha across the seven tests was .75 (min = .65, max = .86).

### Study 2: Selecting the items that best captured variation in *g*: Item Response Theory

With the aim of further reducing our *g* battery and selecting only the best performing items for each test, in a second study (study 2) we administered the seven tests selected in study 1 to an independent sample. We conducted an item response theory (IRT; (72)) analysis to identify items that best capture individual differences in g and estimated their difficulty, discrimination, and information parameters (see Methods). Since one of the main assumptions of IRT is the unidimensionality of the latent construct, we first fitted a principal component analysis (PCA) and a principal component parallel analysis including all 138 items to determine the number of components or **factors** to retain from PCA and examine whether the assumption of unidimensionality held. Although results of the parallel analysis suggested the existence of 3 components,*n* the adjusted eigenvalue for the first component (15.73) was substantially larger than the eigenvalues obtained for the second and third components (2.50 and 1.24, respectively). Further examination of the scree plot obtained from PCA (**Supplementary Figure 1a**) indicated one dimension, which explained 15.3% of the total variance. In addition, when plotting the first three principal components against one another, we found no evidence for multidimensionality (see **Supplementary Figure 1b, 1c and 1d)**. Therefore, we proceeded to perform IRT analysis. Our IRT analysis proceeded in three stages. First, we inspected item information curves for each of the 138 items included in the seven tests. Information curves indicate how informative (reliable) each item is over a particular range of the latent trait. We identified 37 items characterized by horizontal (completely flat) information curves, indicating items that did not discriminate well at any level of the latent trait; these items were removed.

Second, we removed 51 additional items with flat information curves under a threshold of 0.2 or with information curves out of range, either extremely high or low, indicating that the items were either too hard or too easy. This selection process resulted in 20 nonverbal and 30 verbal items.

Third, we focused on refining the verbal battery in order to further reduce the number of items capturing verbal ability. We identified 3 items showing significantly lower information scores than all others; 3 other items with flat item characteristic curves, therefore not discriminating at any level of the latent trait; and 4 additional items that had item characteristic curves that were identical to other items, therefore not providing unique information. These 10 verbal items were deleted, resulting in a battery of 20 nonverbal and 20 verbal items. This 40-item battery included items from all seven tests (**see Supplementary Table 6** for a summary of the reduction process) ranging from very easy (96% of correct responses) to very difficult (7% of correct responses). Information and characteristic curves for the selected items are reported in **Supplementary Figures 2 and 3**, and discrimination and difficulty parameters for all items are reported in **Supplementary Table 7**. Percentages of correct responses and average response times are reported in **Supplementary Table 3**.

These 40 items spanned five tests: 3 verbal ability tests (Vocabulary, Verbal analogies, and Missing letters) and 2 nonverbal ability tests (Matrix reasoning and Visual puzzles). A PCA of the composite scores for the five subdomains showed that the first PC accounted for 67% of the total variance, with factor loadings ranging between .78 and .86 (see **Supplementary Table 8** for the factor loadings).

### Studies 3 and 4: Test-retest reliability and gamification

In studies 3 and 4 we assessed the test-retest reliability of the 40-item measure. In study 3 we examined two-week test-retest reliability, which was excellent for *g*, verbal and nonverbal ability, with phenotypic correlations ranging between .78 (95% CIs= .70, .84) and .89 (95% CIs= .85, .92) (**Supplementary Table 9**). Information on the time limits and order of presentation of each item and subdomain is included in **Supplementary Table 10**.

We proceeded with the process of gamification. Items from each subdomain were embedded into a gamified story line, the *Pathfinder*, which took participants through five ‘journeys’: mountain, tower, woodland, space and ocean (this 2-minute video demonstrates how items were incorporated into the gamified environment: https://www.youtube.com/watch?v=KTk1Ej4F8zE&ab_channel=TEDSProject).

In study 4 we administered *Pathfinder* to the same participants who participated in studies 2 and 3 in order to assess whether the gamification process affected the psychometric properties of the test. **Supplementary Table 10** presents a summary of the Pathfinder journeys and the number and type of items included in each test. Within each sub-domain, items were presented for the same amount of time and in the same order as in study 3. Additional information on *Pathfinder* can be found at the following link: http://www.teds.ac.uk/datadictionary/studies/webtests/21yr_ggame_description.htm.

Study 4, which was conducted approximately 1 month (mean = 29, range = 23 to 35 days) after study 3, showed that test-retest reliability and external validity (i.e., association with education level) remained excellent for *g*, verbal and nonverbal ability even following the gamification of the 40 items .The test-retest correlations ranged between .78 and .91, while the correlations between *g*, verbal and nonverbal ability and education level ranged from .36 to .45 (**Supplementary Table 9**). We also compared the factor structure obtained across the two versions of the test (i.e., study 3 vs. the gamified version administered in study 4) by including the 40 items in two separate CFA model, one for each study. The factor scores derived from each one-factor CFA model correlated at .86, p< .0001, as shown in **Supplementary Figure 4**.

### Study 5: Testing the new *g* measure in a large sample of young adults: Distributions, sex differences, dimensionality and intercorrelations

In study 5, we administered the new *g* measure (*Pathfinder*) to 4,751 twins from the Twins Early Development Study (see Method section and (46) for an in-depth description of the sample). This allowed us to conduct in-depth developmental and genetic analyses to further characterize *Pathfinder*. The first requirement of a good measure of *g* is that it should be distributed normally. We found that the scores for the *g*, verbal and nonverbal ability composites were normally distributed **(see Figure 3A)**. We subsequently investigated sex differences in *g*, verbal ability and nonverbal ability using univariate analysis of variance (ANOVA). Sex differences were significant but small, accounting for between 1 and 3% of the variance. Males outperformed females across the three composites and in four out of five tests, the only exception was performance in the Missing Letter test, for which we found no significant sex differences (**Figure 3B** and **Supplementary Table 11 and 12** for the same analyses in cognitive measures collected over development).

**Figure 3.**
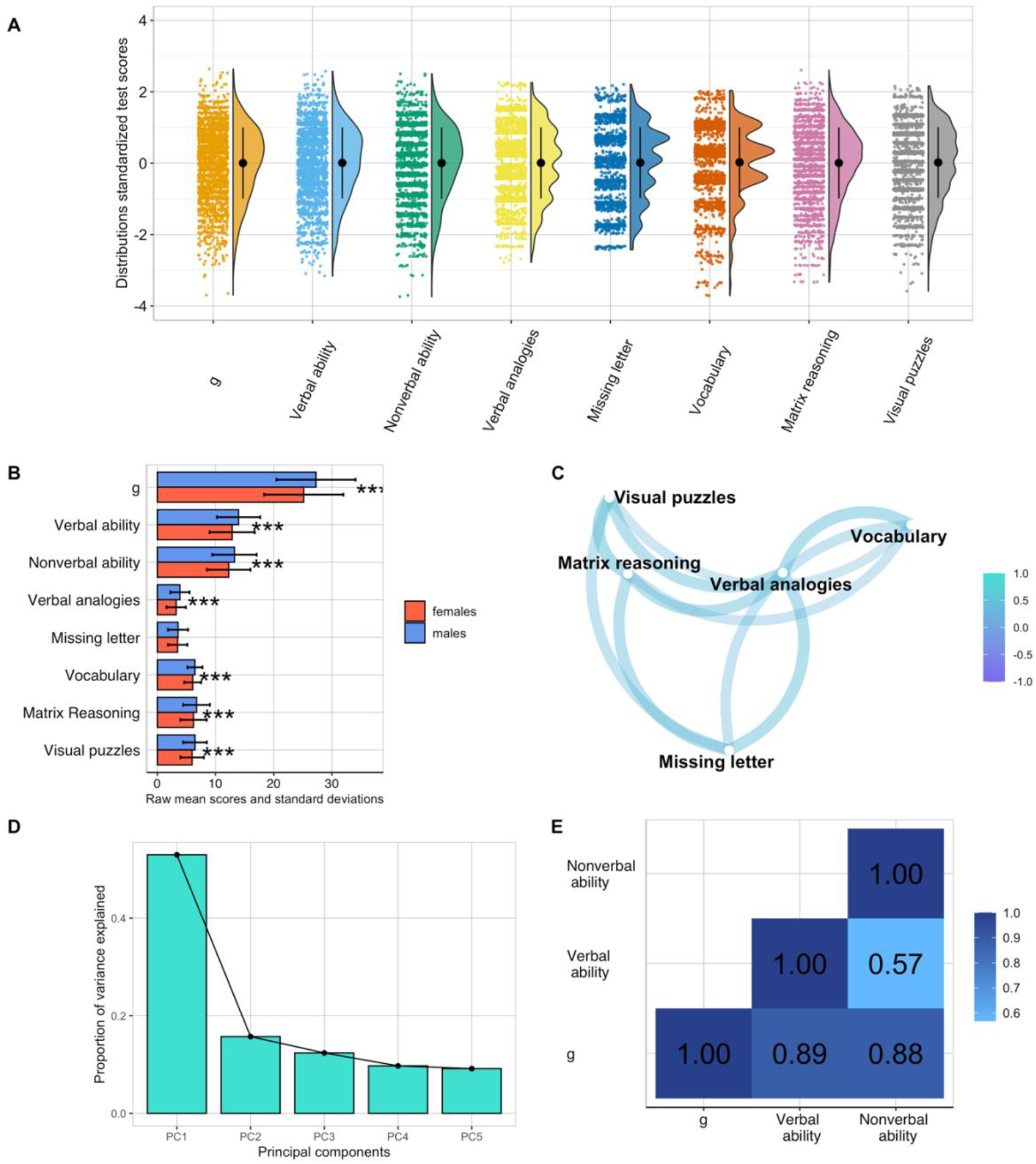
Visual summary of the descriptive properties of the Pathfinder measure administered in study 5. (a) distributions of standardized test scores for the g, verbal and nonverbal ability composites as well as for each subdomain. The colored dots indicate individuals’ performance in each test, black dots represent means and error bars indicate standard deviations for the standardized scores. (b) sex differences in performance across all subdomains and composite scores, *** = p< .001 (two-tailed). (c) Network plot showing the correlations between subdomains, the greater the proximity between points, the greater the correlation between pairs of subdomains. (d) Scree plot of the proportion of variance explained by the principal components. (e) phenotypic correlations between pathfinder composite scores: g, verbal and nonverbal ability.

A second requirement of a good measure of *g* is that it should tap into correlated, yet distinct components of cognitive functioning. We explored this examining the observed correlations between performance in the five subdomains of cognitive ability, which ranged from moderate to strong, as shown in **Figure 3C**. The network plot in **Figure 3C** shows how performance in verbal tests, particularly vocabulary and verbal analogies created a verbal ability cluster, which was correlated with, but more distant from, the nonverbal ability cluster that comprised matrix reasoning and visual puzzles. Performance in the missing letter test was moderately correlated with verbal tests (*r* = .47 and .35 with verbal analogies and vocabulary, respectively) and nonverbal tests (*r* = .43 and .37 with matrix reasoning and visual puzzles, respectively). Correlations between all tests are reported in **Supplementary Table 13**.

A third requirement of a good measure of *g* is that it should produce a first PC accounting for a substantial portion of variance across several cognitive tests, typically about 40%. A PCA of our five tests yielded a first PC that accounted for 52% of variance (**Figure 3D** and **Supplementary Table 14a**). Scores on this first PC correlated .99 with a composite score of *g* created by taking the sum of performance across the 40 verbal and nonverbal items and .99 with a latent factor of *g* created using confirmatory factor analysis (CFA). The results of this CFA analysis are reported in **Supplementary Table 14b**. A one-factor CFA provided a good fit for the data (CFI = 0.95, TLI = 0.90, SRMS = 0.03) and accounted for 73% of the common variance and between 30.1% and 53.1% of the variance in each of the five tests (see **Supplementary Figure 5**). Reliability for this novel *g* measure was high, as indicated by a Cronbach’s alpha of 0.78 and a hierarchical omega coefficient of 0.68.

Considering the nearly perfect correlations between different way of aggregating across cognitive tests, we henceforth consider a composite of *g* constructed from the sum of all items (see Method), a more straightforward approach to compositing. As expected, this *g* composite correlated strongly with verbal ability (.89) and nonverbal ability (.88), while the verbal and nonverbal ability composites correlated with each other to a lesser extent (.57; **Figure 3E**).

### External validity: Performance in *Pathfinder* correlates strongly with cognitive performance measured using well-established cognitive tests, with academic achievement and with family socioeconomic status during childhood and adolescence

Given the developmental nature of the TEDS sample and the rich cognitive and educational data collected from early childhood to emerging adulthood, in study 5 we also examined how well performance in *Pathfinder* mapped onto well-established developmental indicators of cognitive and academic performance assessed at ages 7 to 18. Correlations between the *Pathfinder* composites (*g*, verbal and nonverbal ability) and the corresponding composites created from these other cognitive measures are presented in **Figure 4A-C**. Overall, correlations were strong and increased with age, ranging from .42 at age 7 to .57 at age 16 for *g*, from .39 to .45 for verbal ability and from .28 to .52 for nonverbal ability (**Supplementary Table 15**).

**Figure 4.**
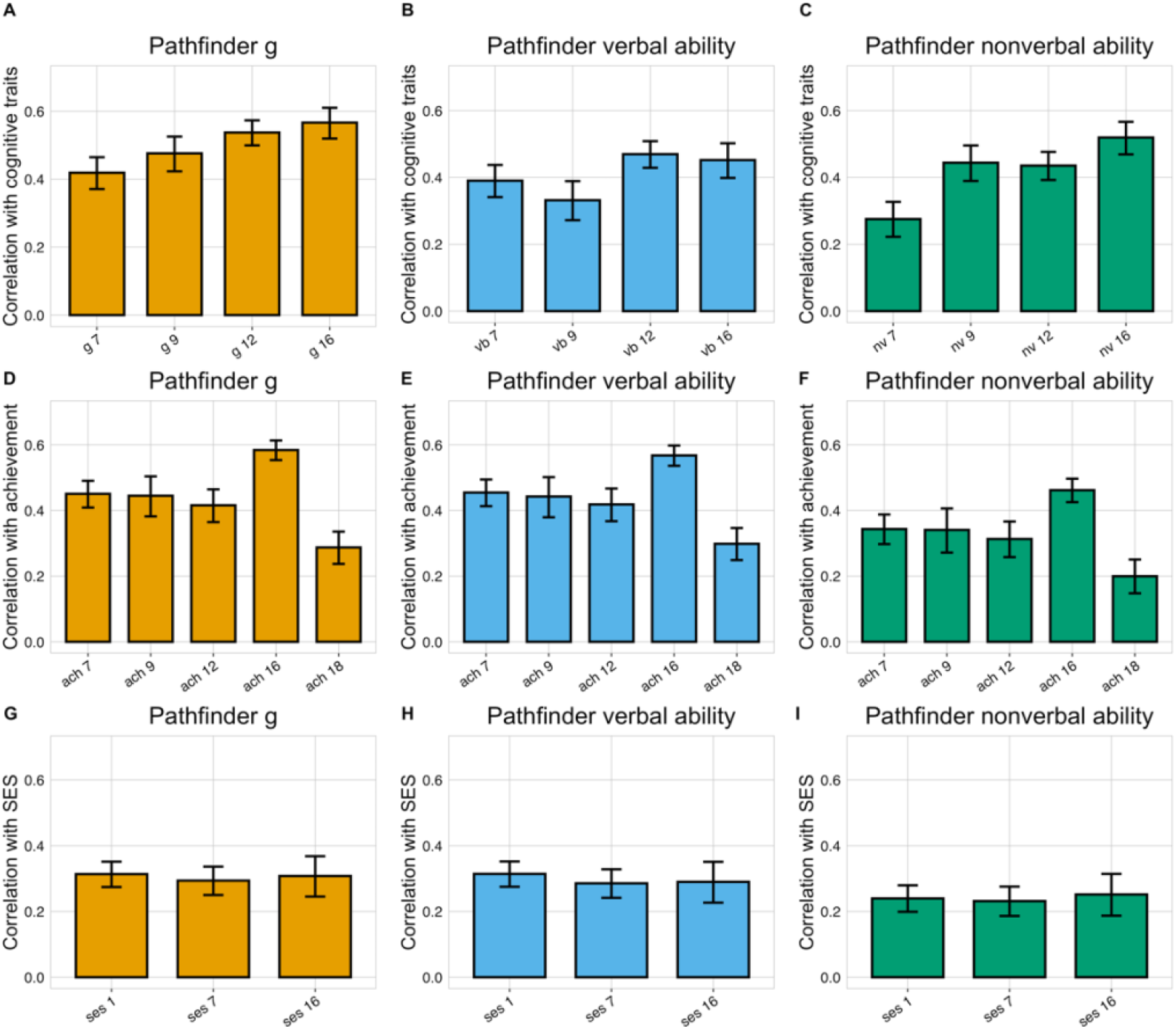
External validity: phenotypic correlations between Pathfinder g, verbal, and nonverbal composites and cognitive (A-C), achievement (D-F) and family socioeconomic status (G-I) measures over development. vb = verbal ability, nv = nonverbal ability, ach = academic achievement, ses = family socioeconomic status. The numbers following each variable name indicate age in years. The length of each bar represents the size of the correlation, and the error bars indicate 95% confidence interval (CIs).

*Pathfinder* composites were also found to be strongly linked to academic achievement during the period of compulsory education, correlations were observed to increase developmentally, ranging from .45 at age 7 to .58 at age 16 for *g*, from .45 to .57 for verbal ability and from .34 to .46 for nonverbal ability (**Figure 4D-F** and **Supplementary Table 15**). The correlation between *Pathfinder* composites and academic performance at age 18, measured with A-level exam grades (see Method) was found to be lower, ranging between .20 and .30, likely due to a restriction of variance as the measure included only those individuals who had continued their education and had taken A-level exams.

In order to further examine how *Pathfinder* related to constructs known to be associated with traditional measures of cognitive ability, we examined the association between the *Pathfinder* composites and family socioeconomic status (SES). In line with the research literature, correlations with SES over development were modest to moderate and similar across the *Pathfinder* composites (average *r* = .30 for *g* and verbal ability and .25 for nonverbal ability; **Figure 4G-I** and **Supplementary Table 15**).

In a further set of analyses, we examined the extrinsic convergent validity (73–75) of Pathfinder relative to other well-known measures of cognitive functioning. Specifically, we compared the external correlational profile of Pathfinder to those of other standardized measures of g as well as verbal and nonverbal ability collected in the TEDS sample over development. These external criteria included measures of academic achievement and SES. **Supplementary Table 16** reports the results of these analyses, which show excellent extrinsic convergent validity for g, verbal, and nonverbal Pathfinder composites. All these measures are functionally equivalent and empirically interchangeable and appear to be indexing the same underlying source of individual difference, general intellectual ability (or *g*).

### *Pathfinder g*, verbal and nonverbal ability show substantial heritability in twin and DNA analyses

A further key requirement for this novel measure was that it should show substantial heritability for two reasons. First, a meta-analysis of cognitive measures across the lifespan yielded an average heritability of 47% (Polderman et al. 2015). Second, substantial heritability is crucial in order for *Pathfinder* to foster genomic discoveries in the cognitive domain. We quantified the heritability of *Pathfinder g*, verbal and nonverbal ability indirectly from the classical twin design and directly from variation in single nucleotide polymorphisms (SNPs) in unrelated individuals (see Method for a description of both techniques).

Twin correlations profiled by zygosity (see **Supplementary Table 17**) revealed substantial differences in MZ and DZ resemblance across the three *Pathfinder* composites: DZ correlation were about half the MZ correlations (**Supplementary Table 17**). In line with the twin correlations, univariate twin model fitting revealed substantial heritability (h^2^) for *Pathfinder g* (h^2^ = .57; 95% CIs = .43, .68), verbal ability (h^2^ = .63; 95% CIs = .49, .69) and nonverbal ability (h^2^ = .46; 95% CIs = .29; .55) and minor shared environmental influences (.08, .03 and .05, respectively) (**Figure 5A**). (See **Supplementary Table 18** for model-fitting estimates and **Supplementary Table 19** for model fit indices). Twin correlations calculated separately for sex and zygosity indicated potential qualitative sex differences (**Supplementary Table 17**) (i.e., differences in same-sex and opposite-sexes DZ twin correlations) for *g* (r = .35 for same sex vs. .25 for opposite sex twins), verbal (.33 vs. .24) and non-verbal (.26 vs. .18) ability. However, formal twin sex-limitation model fitting (**Supplementary Table 20**) showed that both qualitative and quantitative (i.e., differences in MZ-DZ similarity between males and females) sex differences were not significant, indicating that the same genetic effects operate in males and females (76).

**Figure 5.**
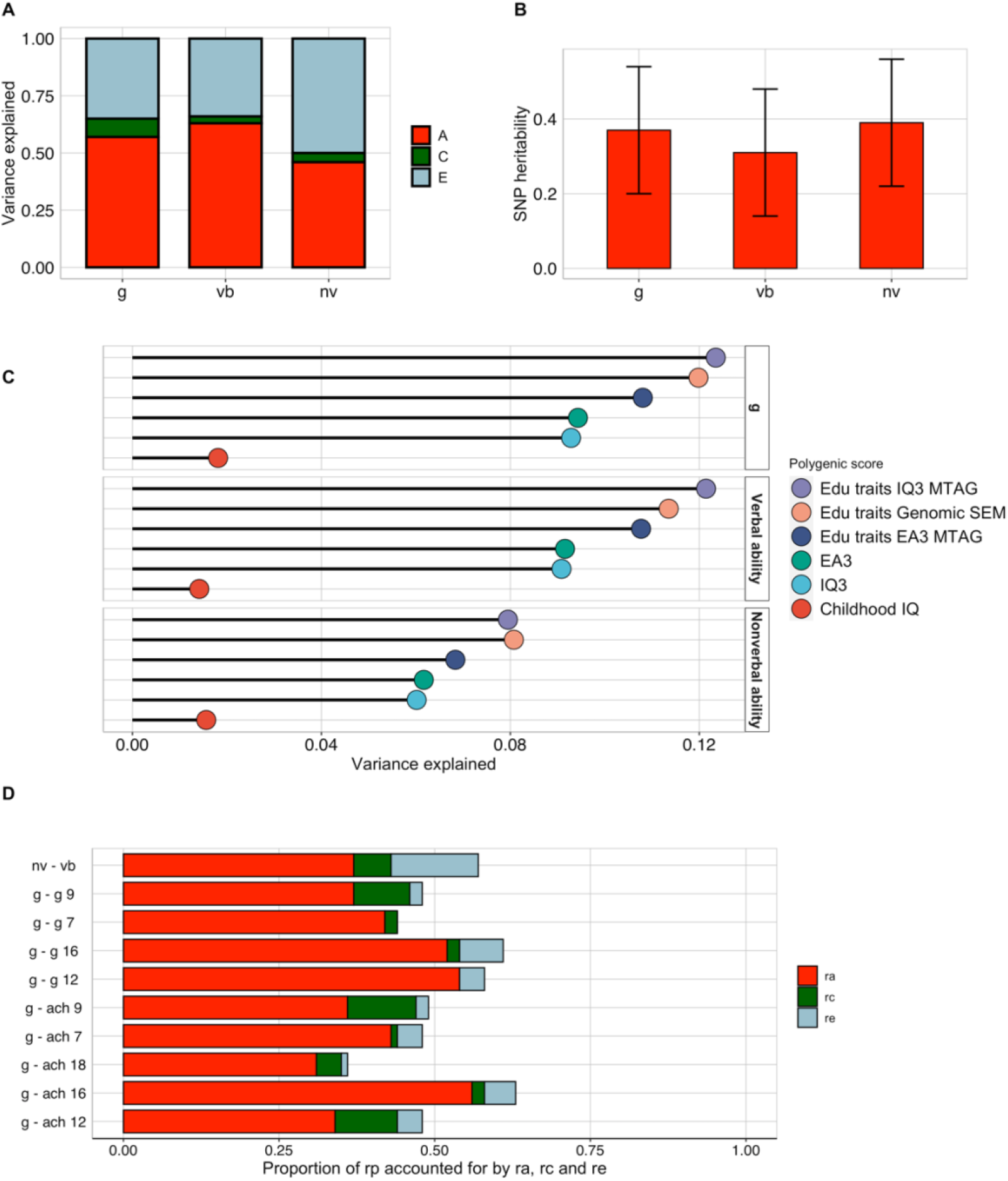
Twin, SNP and polygenic score heritability for Pathfinder composites, and genetic and environmental associations with measures of *g* during childhood and adolescence. **A**. Proportion of variance in Pathfinder g, verbal and nonverbal ability accounted for by heritability, shared environment and nonshared environment calculated using twin design. **B**. SNP heritability estimates (represented by the length of the red bars) and standard errors (represented by the error bars) for Pathfinder *g*, verbal and nonverbal ability composites calculated using GCTA/GREML. **C**. Univariate and multivariate genome-wide polygenic score (GPS) predictions of Pathfinder *g*, verbal and nonverbal ability. **D**. Proportion of the phenotypic correlation between Pathfinder *g* and cognitive and achievement measures accounted for by their genetic (r_A_), shared environmental (r_C_) and nonshared environmental (r_E_) correlation using the twin design. The length of each bar indicates the size of the phenotypic correlation

SNP-based heritability, calculated using GCTA-GREML (see Method), was substantial for the three Pathfinder composites (SNP h^2^ = .37 (SE = .17) for *g*, h^2^ = .31 (SE = .17) for verbal ability and h^2^ = .39 (SE = .17) for nonverbal ability, see **Figure 5B** and **Supplementary Table 21**), around half of the twin heritability estimates. The large standard errors around the estimates indicate that the point estimates were not significantly different, a product of the modest sample size (N = 1,365 unrelated individuals).

We also examined polygenic score heritability: the extent to which genome-wide polygenic scores (GPS, see Method) constructed from GWA studies of cognitive and educationally relevant traits predicted variance in performance in *Pathfinder g*, verbal and nonverbal ability. Specifically, we examined the extent to which the individual GPS based on predictions of childhood IQ (67), adult cognitive performance (IQ3)(23) and educational attainment (EA3)(32) predicted variation in Pathfinder *g*, verbal ability and nonverbal ability. These GPS accounted for between 2% and 9% of the variance in *Pathfinder g*, between 1% and 9% in verbal ability and between 2% and 6% in nonverbal ability (**Figure 5C**, bottom three lines in each case, and **Supplementary Table 22**).

Following examination of how individual GPSs related to variation in performance, we applied multivariate genomic methods to construct GPS aggregating findings from GWAS based on predictions of five cognitive and educationally relevant traits: IQ3, EA3, household income (70), age at completion of full-time education (69) and time spent using computer. Multivariate GPS improved prediction of cognitive measures, accounting for up to 12% of the variance in *Pathfinder g* (β= 0.35, SE = 0.02, t = 19.85, p < .001), up to 12% of the variance in verbal ability (β = 0.35, SE = 0.02, t = 19.67, p <.001) and up to 8% of the variance in nonverbal ability (β = 0.28, SE = 0.02, t = 15.52, p<.001; **Figure 5C**). **Supplementary Table 22** presents these results separately for males and females: GPS prediction were comparable between males and females. This provides support for the potential utility of administering *Pathfinder* to large cohorts to advance our knowledge of the genetics of cognitive ability.

A further characteristic of tests of cognitive ability is that they overlap genetically with many other traits (indexing pleiotropy), and particularly so with other cognitive and educational traits. Genetic correlations (rA) between Pathfinder and other traits were derived from bivariate twin model fitting (see Method). Genetic correlations were substantial between the three *Pathfinder* composites (rA ranging between .73 (95% CIs = .68, .81) and .94 (95% CIs = .92; .96)) and with cognitive and educational measures at earlier ages (rA ranging between .43 (95% CIs = .39, .60) and .95 (95% CIs = .89, 1.00)) (**Supplementary Table 23**). In addition to estimating the extent to which two traits overlap genetically, bivariate twin model fitting also estimates the extent to which they overlap for environmental reasons. Shared environmental correlations, indicating how similarities between family members contribute to the association between traits, were mostly not significant. On the other hand, nonshared environmental correlations, pointing to how environmental experiences that differ between siblings contribute to the association between two traits, were modest between *Pathfinder* composites (rE = .33; 95% CIs = .26, .39) but small with cognitive and educational measures obtained at earlier ages, with rE ranging between -0.03 (95% CIs = -.12, .07) and 0.28 (95% CIs = .18, .37) (**Supplementary Table 23**).

Bivariate associations between traits can also be expressed in terms of the proportion of their phenotypic correlations that is accounted for by genetic, shared environmental and nonshared environmental factors, respectively. For example, genetic factors accounted for 64.9% of the correlation between *Pathfinder* verbal ability and nonverbal ability, shared environmental factors accounted for 10.5% of their correlation and nonshared environmental factors accounted for 24.6% of their correlation. (**Figure 5D**, with fit statistics in **Supplementary Table 24**). **Figure 5D** also shows the proportional contribution of genetics (A), shared environment (C) and nonshared environment (E) to the phenotypic correlation between *Pathfinder g* and cognitive performance over development. Estimates for verbal and nonverbal composites are reported in **Supplementary Table 25**.

## Discussion

*Pathfinder* is a 15-minute gamified online test whose construction was guided by item response theory and principal component analysis to be a maximally efficient and reliable measure of *g*. The first principal component accounts for 52% of the total variance, which reflects the communalities among the five tests. The *g* score is normally distributed and its one-month test-retest reliability is .88. Despite the strong *g* factor, we were able to differentiate verbal and non-verbal cognitive abilities, which correlated .57 and yielded one-month test-retest reliabilities of .90 for verbal and .75 for nonverbal. This engaging, freely available and easily accessible measure is a fundamental resource that enables scientists easily to incorporate general cognitive ability in research across the biological, medical, and behavioural sciences.

We were especially interested in the application of *Pathfinder* in genetic studies. In the midst of a replication crisis in science (77), it is noteworthy that genetic and genomic results replicate reliably (78). On the basis of previous research, we predicted (https://osf.io/pc9yh/) that twin heritability for *g* would be greater than 50%, that shared environmental influence would be less than 20% and that multivariate polygenic scores would predict more than 10% of the variance. Our results confirmed these hypotheses: Heritability was 57%, shared environmental influence was 8% and multivariate polygenic scores predicted up to 12% of the variance.

The latter finding – that 12% of the variance of *Pathfinder g* can be predicted by DNA – makes this the strongest polygenic score predictor of *g* reported to date (13). Although 12% is only one fifth of the twin study estimate of heritability, we hope that adding *Pathfinder g* in large biobanks will improve the yield of meta-analytic GWAS analyses by increasing sample sizes and decreasing heterogeneity of cognitive measures. It should be possible to use the brute force method of increasing sample sizes, especially with less heterogeneity of measures, to close the missing heritability gap from 12% to the SNP heritability of about 30%.

A more daunting challenge is to break through the ceiling of 30% SNP heritability to reach the 60% heritability estimated by twin studies of adults. Both GPS heritability and SNP heritability are limited to the additive effects of the common SNPs assessed on SNP chips used in GWAS studies. Going beyond SNP heritability will require whole-genome sequencing that can assess rare variants and methodologies to analyze gene-gene and gene-environment interactions (13).

Nonetheless, predicting 12% of the variance of *g* is a notable achievement for two reasons. First, until 2016 polygenic scores could predict only 1% of the variance in general cognitive ability (13). Predicting a substantial amount of variance (more than 10% in this case) is an important milestone for genetic research on intelligence because effect sizes of this magnitude are large enough to be ‘perceptible to the naked eye of a reasonably sensitive observer’ (79). Second, effect sizes like this, are rare in the behavioural sciences. For example, one of the most widely used predictors of children’s *g* and educational achievement is family SES. We showed that family SES predicts 9% of the variance of *Pathfinder*-assessed *g*. At 2 years of age, infant intelligence tests predict less than 5% of the variance of *g* in late adolescence (80,81). It is not until the early school years that children’s cognitive test scores predict more than 10% of the variance of adult *g*. The unique value of polygenic scores is that their prediction of adult *g* is just as strong from early in life as it is in adulthood because inherited DNA differences do not change. Increasing the predictive power of polygenic scores also opens important new avenues for investigating the mechanisms underlying this prediction, including the environmental experiences that mediate this pathway from genotype to phenotype (24).

We were primarily motivated to create a measure of *g* that could be used in large biobanks to improve the power of meta-analytic GWA studies to identify the minuscule SNP associations we now know to be responsible for the heritability of *g*. However, because *g* pervades so many aspects of life – education, occupation, wealth, and health – we hope that *Pathfinder* will open new avenues for research into the causes and consequences of general cognitive ability throughout the life sciences. Incorporating *g* in biological, medical, and behavioural research can add a new dimension that capitalizes on the pleiotropic power of *g*. Using *Pathfinder* as a standard measure of *g* will also improve the reproducibility of research in the life sciences, which is critical in light of the replication crisis (82). For these reasons, we have designed a platform to make it easy to use Pathfinder. Further information on how to access *Pathfinder* can be found at the following webpage, specifically created for the purpose of sharing the test: www.pathfindertestgame.com

Limitations of the present study point the way to future research. Like most genetic and genomic research, the results of our study cannot be safely generalized beyond its UK sample whose ancestry is 90% northern European. Although twin study heritability estimates of *g* are substantial in other countries and ancestries (83,84), polygenic scores derived largely from GWAS of northern European samples are not yet as predictive in other ancestral groups (85). The present study has three more practical limitations. First, *Pathfinder* is as yet limited to English, although the test’s language load is light, which will render translation, including appropriate linguistic and cultural adaptation, manageable. Second, no alternate forms have as yet been created, which would be useful for longitudinal designs that require repeated testing, although the high one-month test-retest reliability suggests that the *Pathfinder* test can be used for repeated testing. Third, *Pathfinder* was created in samples of adults from 18 to 49 years of age, so its utility for younger or older groups remains to be investigated.

One of the most widely adopted definition of *g* describes it as “…a very general mental capability that, among other things, involves the ability to reason, plan, solve problems, think abstractly, comprehend complex ideas, learn quickly and learn from experience.” (Gottfredson, 1997, p. 13). Alternative conceptualizations and interpretations have also been proposed, most notably the view that *g* does not reflect a set of a domain general abilities, but is in fact mental energy (86), or a property of the mind (87), potentially simply indexing overall cognitive potential. However the statistical abstraction of *g* is interpreted, its remarkable ability to predict important functional and life outcomes, and its likely universality supported by cross-cultural research (88,89), a deeper understanding of *g* has the potential to lead to major scientific advances in our understanding of human development from several scientific angles, from molecular genetics to psychology and evolutionary biology.

To conclude, over four studies we have created a very brief (15-minute), reliable and valid measure of *g, Pathfinder*, that given its gamified features, is also engaging. *Pathfinder* can be accessed by all researchers, and easily integrated within existing data collection platforms. It is our hope that widespread use of this engaging new measure will advance research not only in genomics but throughout the biological, medical, and behavioural sciences.

## Acknowledgements

We gratefully acknowledge the ongoing contribution of the participants in the Twins Early Development Study (TEDS) and their families. TEDS is supported by a program grant to RP from the UK Medical Research Council (MR/M021475/1 and previously G0901245), with additional support from the US National Institutes of Health (AG046938). RP is supported by a Medical Research Council Professorship award (G19/2). KR is supported by a Sir Henry Wellcome Postdoctoral Fellowship. This study presents independent research [part-] funded by the National Institute for Health Research (NIHR) Biomedical Research Centre at South London and Maudsley NHS Foundation Trust and King’s College London. AG is supported by a Queen Mary School of Biological and Chemical Sciences PhD Fellowship awarded to MM. We thank Emily Smith-Woolley and Ziada Ayorech for their help with the initial phases of this work.

## Code availability

Code will be made available by the corresponding author upon request.

## References

1. Deary IJ. Intelligence. Annu Rev Psychol [Internet]. 2011 Nov 30;63(1):453–82. Available from: https://doi.org/10.1146/annurev-psych-120710-100353

2. Spearman C. “ General Intelligence, ” Objectively Determined and Measured Author (s): C. Spearman Source : The American Journal of Psychology, Vol. 15, No. 2 (Apr., 1904), pp. 201-292 Published by : University of Illinois Press Stable URL : http://www.jsto. Am J Psychol. 1904;15(2):201–92.

3. Gottfredson LS. Why g matters: The complexity of everyday life. Intelligence [Internet]. 1997;24(1):79–132. Available from: https://www.sciencedirect.com/science/article/pii/S0160289697900143

4. Gottfredson LS. Intelligence: Is It the Epidemiologists’ Elusive “Fundamental Cause” of Social Class Inequalities in Health? Vol. 86, Journal of Personality and Social Psychology. Gottfredson, Linda S.: School of Education, University of Delaware, Willard Hall, Newark, DE, US, 19716-2922,: American Psychological Association; 2004. p. 174–99.

5. Carroll JB (John B. Human cognitive abilities : a survey of factor-analytic studies. Cambridge University Press; 1993. 819 p.

6. Chipuer HM, Rovine MJ, Plomin R. LISREL modeling: Genetic and environmental influences on IQ revisited. Intelligence. 1990;14(1):11–29.

7. Tucker-Drob EM, Briley DA. Continuity of genetic and environmental influences on cognition across the life span: A meta-analysis of longitudinal twin and adoption studies. Psychol Bull. 2014;140(4):949–79.

8. Haworth CMA, Wright MJ, Luciano M, Martin NG, De Geus EJC, Van Beijsterveldt CEM, et al. The heritability of general cognitive ability increases linearly from childhood to young adulthood. Mol Psychiatry [Internet]. 2010;15(11):1112–20. Available from: http://dx.doi.org/10.1038/mp.2009.55

9. Plomin R, Fulker DW, Corley R, DeFries JC. Nature, Nurture, and Cognitive Development from 1 to 16 Years: A Parent-Offspring Adoption Study. Psychol Sci [Internet]. 1997 Nov 1;8(6):442–7. Available from: https://doi.org/10.1111/j.1467-9280.1997.tb00458.x

10. Petrill SA. The case for general intelligence: A behavioral genetic perspective. In: The general factor of intelligence: How general is it? Mahwah, NJ, US: Lawrence Erlbaum Associates Publishers; 2002. p. 281–98.

11. Plomin R, Kovas Y. Generalist Genes and Learning Disabilities. Psychol Bull [Internet]. 2005;131(4):592–617. Available from: http://doi.apa.org/getdoi.cfm?doi=10.1037/0033-2909.131.4.592

12. de la Fuente J, Davies G, Grotzinger AD, Tucker-Drob EM, Deary IJ. A general dimension of genetic sharing across diverse cognitive traits inferred from molecular data. Nat Hum Behav [Internet]. 2020; Available from: https://doi.org/10.1038/s41562-020-00936-2

13. Plomin R, Von Stumm S. The new genetics of intelligence. Nat Rev Genet. 2018;19(3):148–59.

14. Davies NM, Hill WD, Anderson EL, Sanderson E, Deary IJ, Davey Smith G. Multivariable two-sample Mendelian randomization estimates of the effects of intelligence and education on health. Teare MD, Franco E, Burgess S, editors. Elife [Internet]. 2019;8:e43990. Available from: https://doi.org/10.7554/eLife.43990

15. Daly M, Egan M, O’Reilly F. Childhood general cognitive ability predicts leadership role occupancy across life: Evidence from 17,000 cohort study participants. Leadersh Q [Internet]. 2015;26(3):323– 41. Available from: http://dx.doi.org/10.1016/j.leaqua.2015.03.006

16. Kalechstein AD, Newton TF, van Gorp WG. Neurocognitive Functioning is Associated ith Employment Status: A Quantitative Review. J Clin Exp Neuropsychol (Neuropsychology, Dev Cogn Sect A) [Internet]. 2003;25(8):1186–91. Available from: http://www.tandfonline.com/doi/abs/10.1076/jcen.25.8.1186.16723

17. Mollon J, David AS, Zammit S, Lewis G, Reichenberg A. Course of Cognitive Development From Infancy to Early Adulthood in the Psychosis Spectrum. JAMA Psychiatry [Internet]. 2018;06510. Available from: http://archpsyc.jamanetwork.com/article.aspx?doi=10.1001/jamapsychiatry.2017.4327

18. Snyder HR, Miyake A, Hankin BL. Advancing understanding of executive function impairments and psychopathology: bridging the gap between clinical and cognitive approaches. Front Psychol [Internet]. 2015 Mar 26;6. Available from: http://www.frontiersin.org/Psychopathology/10.3389/fpsyg.2015.00328/abstract

19. Latvala A, Kuja-Halkola R, D’Onofrio BM, Larsson H, Lichtenstein P. Cognitive ability and risk for substance misuse in men: genetic and environmental correlations in a longitudinal nation-wide family study. Addiction. 2016;111(10):1814–22.

20. Deary IJ, Weiss A, Batty GD. Intelligence and personality as predictors of illness and death: How researchers in differential psychology and chronic disease epidemiology are collaborating to understand and address health inequalities. Psychol Sci Public Interes Suppl. 2010;11(2):53–79.

21. Smeland OB, Bahrami S, Frei O, Shadrin A, O’Connell K, Savage J, et al. Genome-wide analysis reveals extensive genetic overlap between schizophrenia, bipolar disorder, and intelligence. Mol Psychiatry [Internet]. 2020;25(4):844–53. Available from: https://doi.org/10.1038/s41380-018-0332-x

22. Marioni RE, Davies G, Hayward C, Liewald D, Kerr SM, Campbell A, et al. Molecular genetic contributions to socioeconomic status and intelligence. Intelligence [Internet]. 2014;44:26–32. Available from: http://www.sciencedirect.com/science/article/pii/S0160289614000178

23. Savage JE, Jansen PR, Stringer S, Watanabe K, Bryois J, Leeuw CA de, et al. GWAS meta-analysis (N=279,930) identifies new genes and functional links to intelligence. Nat Genet [Internet]. 2018;50:912–9. Available from: http://www.pnas.org/lookup/doi/10.1073/pnas.1605859113

24. Malanchini M, Rimfeld K, Allegrini AG, Ritchie SJ, Plomin R. Cognitive ability and education: How behavioural genetic research has advanced our knowledge and understanding of their association. Neurosci Biobehav Rev [Internet]. 2020;111(July 2019):229–45. Available from: https://doi.org/10.1016/j.neubiorev.2020.01.016

25. Demange PA, Malanchini M, Mallard TT, Biroli P, Cox SR, Grotzinger AD, et al. Investigating the Genetic Architecture of Non-Cognitive Skills Using GWAS-by-Subtraction. bioRxiv. 2020;2020.01.14.905794.

26. Ormel J, Hartman CA, Snieder H. The genetics of depression: successful genome-wide association studies introduce new challenges. Transl Psychiatry [Internet]. 2019;9(1):114. Available from: https://doi.org/10.1038/s41398-019-0450-5

27. Ripke S, Walters JTR, O’Donovan MC. Mapping genomic loci prioritises genes and implicates synaptic biology in schizophrenia. medRxiv [Internet]. 2020 Jan 1;2020.09.12.20192922. Available from: http://medrxiv.org/content/early/2020/09/13/2020.09.12.20192922.abstract

28. Howard DM, Adams MJ, Clarke TK, Hafferty JD, Gibson J, Shirali M, et al. Genome-wide meta-analysis of depression identifies 102 independent variants and highlights the importance of the prefrontal brain regions. Nat Neurosci [Internet]. 2019;22(3):343–52. Available from: http://dx.doi.org/10.1038/s41593-018-0326-7

29. Mullins N, Forstner AJ, O’Connell KS, Coombes B, Coleman JRI, Qiao Z, et al. Genome-wide association study of over 40,000 bipolar disorder cases provides novel biological insights. medRxiv [Internet]. 2020 Jan 1;2020.09.17.20187054. Available from: http://medrxiv.org/content/early/2020/09/18/2020.09.17.20187054.abstract

30. Allegrini AG, Selzam S, Rimfeld K, von Stumm S, Pingault JB, Plomin R. Genomic prediction of cognitive traits in childhood and adolescence. Mol Psychiatry [Internet]. 2019;000:819–27. Available from: http://dx.doi.org/10.1038/s41380-019-0394-4

31. Chabris CF, Lee JJ, Cesarini D, Benjamin DJ, Laibson DI. The Fourth Law of Behavior Genetics. Curr Dir Psychol Sci. 2015;24(4):304–12.

32. Lee JJ, Wedow R, Okbay A, Kong E, Maghzian O, Zacher M, et al. Gene discovery and polygenic prediction from a genome-wide association study of educational attainment in 1.1 million individuals. Nat Genet. 2018;50(8):1112–21.

33. Génin E. Missing heritability of complex diseases: case solved? Hum Genet [Internet]. 2020;139(1):103–13. Available from: https://doi.org/10.1007/s00439-019-02034-4

34. Wainschtein P, Jain DP, Yengo L, Zheng Z, Cupples LA, Shadyab AH, et al. Recovery of trait heritability from whole genome sequence data. bioRxiv [Internet]. 2019 Jan 1;588020. Available from: http://biorxiv.org/content/early/2019/03/25/588020.abstract

35. Nunes A, Trappenberg T, Alda M. The definition and measurement of heterogeneity. Transl Psychiatry [Internet]. 2020;10(1):299. Available from: https://doi.org/10.1038/s41398-020-00986-0

36. Manchia M, Cullis J, Turecki G, Rouleau GA, Uher R, Alda M. The Impact of Phenotypic and Genetic Heterogeneity on Results of Genome Wide Association Studies of Complex Diseases. PLoS One [Internet]. 2013 Oct 11;8(10):e76295. Available from: https://doi.org/10.1371/journal.pone.0076295

37. Cai N, Revez JA, Adams MJ, Andlauer TFM, Breen G, Byrne EM, et al. Minimal phenotyping yields genome-wide association signals of low specificity for major depression. Nat Genet [Internet]. 2020;52(4):437–47. Available from: https://doi.org/10.1038/s41588-020-0594-5

38. de Vlaming R, Okbay A, Rietveld CA, Johannesson M, Magnusson PKE, Uitterlinden AG, et al. Meta-GWAS Accuracy and Power (MetaGAP) Calculator Shows that Hiding Heritability Is Partially Due to Imperfect Genetic Correlations across Studies. PLOS Genet [Internet]. 2017 Jan 17;13(1):e1006495. Available from: https://doi.org/10.1371/journal.pgen.1006495

39. Zainuddin Z, Shujahat M, Haruna H, Chu SKW. The role of gamified e-quizzes on student learning and engagement: An interactive gamification solution for a formative assessment system. Comput Educ [Internet]. 2020;145:103729. Available from: http://www.sciencedirect.com/science/article/pii/S0360131519302829

40. Buckley P, Doyle E. Gamification and student motivation. Interact Learn Environ [Internet]. 2016 Aug 17;24(6):1162–75. Available from: https://doi.org/10.1080/10494820.2014.964263

41. Pike GR, Graunke SS. Examining the Effects of Institutional and Cohort Characteristics on Retention Rates. Res High Educ [Internet]. 2015;56(2):146–65. Available from: http://www.jstor.org/stable/24572009

42. Sudlow C, Gallacher J, Allen N, Beral V, Burton P, Danesh J, et al. UK Biobank: An Open Access Resource for Identifying the Causes of a Wide Range of Complex Diseases of Middle and Old Age. PLOS Med [Internet]. 2015 Mar 31;12(3):e1001779.

43. Hampshire A, Trender W, Chamberlain SR, Jolly A, Grant JE, Patrick F, et al. Cognitive deficits in people who have recovered from COVID-19 relative to controls: An N=84,285 online study. medRxiv [Internet]. 2020 Jan 1

44. Fawns-Ritchie C, Deary IJ. Reliability and validity of the UK Biobank cognitive tests. PLoS One [Internet]. 2020 Apr 20;15(4):e0231627.

45. Rimfeld K, Malanchini M, Spargo T, Spickernell G, Selzam S, McMillan A, et al. Twins Early Development Study: A Genetically Sensitive Investigation into Behavioral and Cognitive Development from Infancy to Emerging Adulthood. Twin Res Hum Genet [Internet]. 2019 Sep 23 [cited 2020 Jan 2];1–6.

46. Rimfeld K, Malanchini M, Spargo T, Spickernell G, Selzam S, McMillan A, et al. Twins Early Development Study: A Genetically Sensitive Investigation into Behavioral and Cognitive Development from Infancy to Emerging Adulthood. Twin Res Hum Genet. 2019;22(6).

47. Raven JC., Raven J., Court J. Mill Hill Vocabulary Scale. Oxford: Oxford: OOP; 1998.

48. Raven JC., Court JH., Raven J. Manual for Raven’s progressive matrices and vocabulary scales. Oxford: Oxford University Press; 1996.

49. Selzam S, McAdams TA, Coleman JRI, Carnell S, O’Reilly PF, Plomin R, et al. Evidence for gene-environment correlation in child feeding: Links between common genetic variation for BMI in children and parental feeding practices. PLoS Genet. 2018;14(11):1–19.

50. Wechsler D. Wechsler Intelligence Scale for Children (3rd Ed. UK). The Psychological Corporation; 1992.

51. McCarthy D. McCarthy Scales of Children’s Abilities. New York: The Psychological Corporation.; 1972.

52. Kaplan E., Fein D., Kramer J., Delis D., Morris R. WISC-III as a process instrument (WISC-III-PI). The Psychological Corporation; 1999.

53. Smith P, Fernandes C, Strand S. Cognitive ability test 3 (CAT3). Windsor, Engalnd: nferNelson; 2001.

54. Raven J, Raven JC, Court J. Manual for Raven’s Progressive Matrices and Vocabulary Scales. Raven manual. Oxford: Oxford University Press; 1996.

55. Shakeshaft NG, Trzaskowski M, McMillan A, Rimfeld K, Krapohl E, Haworth CMA, et al. Strong genetic influence on a UK nationwide test of educational achievement at the end of compulsory education at age 16. PLoS One. 2013;8(12).

56. Muthén LK, Muthén BO. Mplus User’s Guide. J Am Geriatr Soc [Internet]. 2007;2006:676.

57. van den Berg SM, Glas CAW, Boomsma DI. Variance decomposition using an IRT measurement model. Vol. 37, Behavior Genetics. 2007. p. 604–16.

58. Kendler KS, Neale MC, Kessler RC, Heath AC, Eaves L. A twin study of recent life events and difficulties. Arch Gen Psychiatry. 1993;50:789–96.

59. Martin NG, Eaves LJ. Stages ; the First To Determine the Genetical and Environmental Model. Most. 1977;38:79–95.

60. Neale MC, Hunter MD, Pritikin JN, Zahery M, Brick TR, Kirkpatrick RM, et al. OpenMx 2.0: Extended Structural Equation and Statistical Modeling. Psychometrika [Internet]. 2016;81(2):535–49.

61. Rijsdijk F V, Sham PC. Analytic approaches to twin data using structural equation models. Brief Bioinform [Internet]. 2002;3(2):119–33.

62. Yang J, Lee SH, Goddard ME, Visscher PM. GCTA: A tool for genome-wide complex trait analysis. Am J Hum Genet [Internet]. 2011;88(1):76–82. Available from: http://dx.doi.org/10.1016/j.ajhg.2010.11.011

63. Yang J, Benyamin B, McEvoy BP, Gordon S, Henders AK, Nyholt DR, et al. Common SNPs explain a large proportion of the heritability for human height. Nat Genet. 2010;42(7):565–9.

64. Vilhjálmsson BJ, Yang J, Finucane HK, Gusev A, Lindström S, Ripke S, et al. Modeling linkage disequilibrium increases accuracy of polygenic risk scores. Am J Hum Genet. 2015;97(4):576–92.

65. Allegrini AG, Selzam S, Rimfeld K, von Stumm S, Pingault JB, Plomin R. Genomic prediction of cognitive traits in childhood and adolescence. Mol Psychiatry. 2019;1–24.

66. Lee JJ, Wedow R, Okbay A, Kong E, Maghzian O, Zacher M, et al. Gene discovery and polygenic prediction from a genome-wide association study of educational attainment in 1.1 million individuals. Nat Genet. 2018;50(8):1112–21.

67. Benyamin B, Pourcain Bs, Davis OS, Davies G, Hansell NK, Brion M-J, et al. Childhood intelligence is heritable, highly polygenic and associated with FNBP1L. Mol Psychiatry [Internet]. 2014;19(2):253–8. Available from: https://doi.org/10.1038/mp.2012.184

68. Turley P, Walters RK, Maghzian O, Okbay A, Lee JJ, Fontana MA, et al. Multi-trait analysis of genome-wide association summary statistics using MTAG. Nat Genet. 2018;50(2):229–37.

69. Grotzinger AD, Rhemtulla M, de Vlaming R, Ritchie SJ, Mallard TT, Hill WD, et al. Genomic structural equation modelling provides insights into the multivariate genetic architecture of complex traits. Nat Hum Behav [Internet]. 2019;3(5):513–25. Available from: https://doi.org/10.1038/s41562-019-0566-x

70. Hill WD, Hagenaars SP, Marioni RE, Harris SE, Liewald DCM, Davies G, et al. Molecular Genetic Contributions to Social Deprivation and Household Income in UK Biobank. Curr Biol [Internet]. 2016 Nov 21;26(22):3083–9. Available from: https://doi.org/10.1016/j.cub.2016.09.035

71. Seed CH. An Open-Source Framework for Scalable Genetic Data. [Internet]. 2017. Available from: http://www.nealelab.is/uk-biobank

72. Reise SP, Ainsworth AT, Haviland MG. Item Response Theory: Fundamentals, Applications, and Promise in Psychological Research. Curr Dir Psychol Sci [Internet]. 2005 Apr 1;14(2):95–101. Available from: https://doi.org/10.1111/j.0963-7214.2005.00342.x

73. Lubinski D. Introduction to the Special Section on Cognitive Abilities: 100 Years After Spearman’s (1904) “‘General Intelligence,’ Objectively Determined and Measured.” J Pers Soc Psychol. 2004;86(1):96–111.

74. Lubinski D, Tellegen A, Butcher JN. Masculinity, femininity, and androgyny viewed and assessed as distinct concepts. J Pers Soc Psychol. 1983;44(2):428–39.

75. Gonzalez O, MacKinnon DP, Muniz FB. Extrinsic convergent validity evidence to prevent jingle and jangle fallacies. Multivariate Behav Res. 2021;56(1):3–19.

76. Vink JM, Bartels M, van Beijsterveldt TCEM, van Dongen J, van Beek JHDA, Distel MA, et al. Sex Differences in Genetic Architecture of Complex Phenotypes? PLoS One [Internet]. 2012 Dec 18;7(12):e47371. Available from: https://doi.org/10.1371/journal.pone.0047371

77. Ritchie SJ. Science Fiction. Penguin; 2020. 368 p.

78. Plomin R, DeFries JC, Knopik VS, Neiderhiser JM. Top 10 Replicated Findings From Behavioral Genetics. Perspect Psychol Sci [Internet]. 2016 Jan 1;11(1):3–23. Available from: https://doi.org/10.1177/1745691615617439

79. Cohen J. Statistical power analysis for the behavioral sciences, Rev. ed. Statistical power analysis for the behavioral sciences, Rev. ed. Hillsdale, NJ, US: Lawrence Erlbaum Associates, Inc; 1977. p. xv, 474–xv, 474.

80. Honzik MP, Macfarlane JW, Allen L. The Stability of Mental Test Performance Between two and Eighteen Years. J Exp Educ [Internet]. 1948 Dec 1;17(2):309–24. Available from: https://doi.org/10.1080/00220973.1948.11010388

81. von Stumm S, Plomin R. Socioeconomic status and the growth of intelligence from infancy through adolescence. Intelligence [Internet]. 2015;48:30–6. Available from: http://dx.doi.org/10.1016/j.intell.2014.10.002

82. Schnack H. Assessing reproducibility in association studies. Elife [Internet]. 2019;8:e46757. Available from: https://doi.org/10.7554/eLife.46757

83. Polderman TJC, Benyamin B, De Leeuw CA, Sullivan PF, Van Bochoven A, Visscher PM, et al. Meta-analysis of the heritability of human traits based on fifty years of twin studies. Nat Genet [Internet]. 2015;47(7):702–9. Available from: http://dx.doi.org/10.1038/ng.3285

84. Knopik VS, Neiderhiser JM, Defries JC, Plomin R. Behavioral Genetics. Seventh. Macmillan Higher Education; 2016.

85. Martin AR, Kanai M, Kamatani Y, Okada Y, Neale BM, Daly MJ. Clinical use of current polygenic risk scores may exacerbate health disparities. Nat Genet [Internet]. 2019;51(4):584–91. Available from: https://doi.org/10.1038/s41588-019-0379-x

86. Spearman C. The abilities of man: Their nature and measurement. Oxford, England: Macmillan; 1927.

87. Jensen AR. The g factor: The science of mental ability. The g factor: The science of mental ability. Westport, CT, US: Praeger Publishers/Greenwood Publishing Group; 1998. xiv, 648–xiv, 648. (Human evolution, behavior, and intelligence.).

88. Warne RT, Burningham C. Spearman’s g found in 31 non-Western nations: Strong evidence that g is a universal phenomenon. Vol. 145, Psychological Bulletin. 2019. p. 237–72.

89. Flaim M, Blaisdell AP. The comparative analysis of intelligence. Psychol Bull. 2020;146(12):1174– 99.

