## Supplementary Figures for "Pathfinder: A gamified measure to integrate general cognitive ability into the biological, medical, and behavioural sciences"

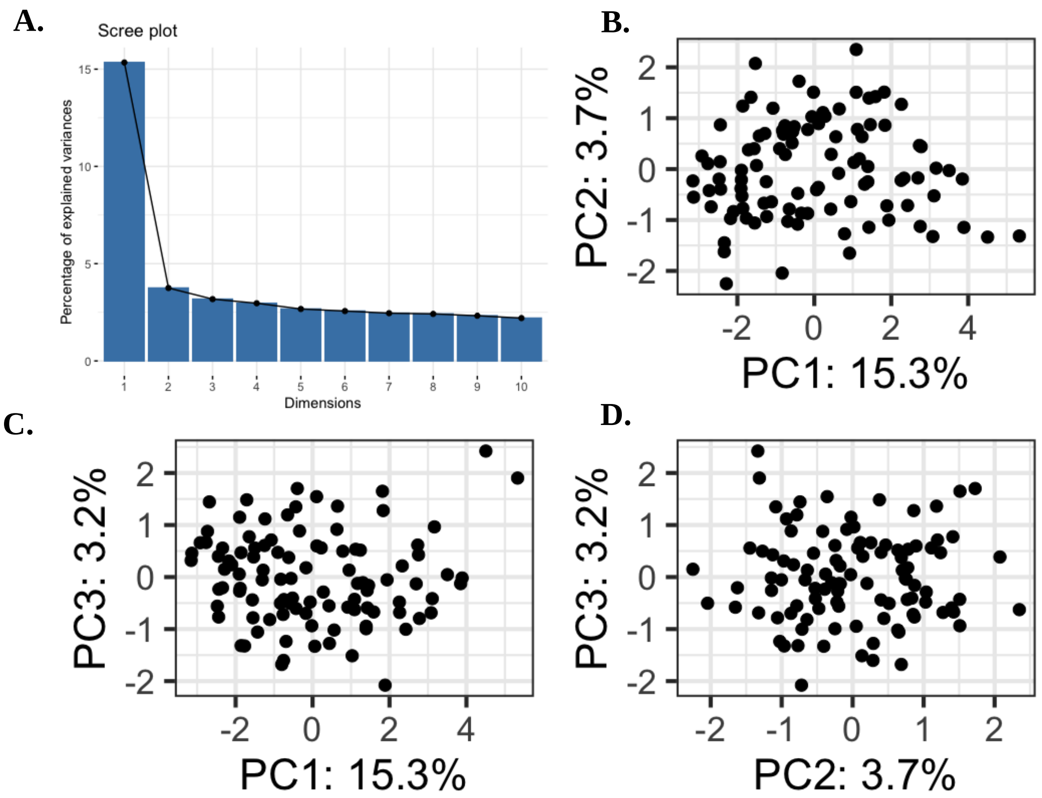


**Supplementary Figure 1**. Testing the dimensionality of the 138 items included in Study 2 prior to IRT analyses. Panel A. shows the scree plot indicating the existence of one main dimension that accounts for 15.3% of the variance. Panel B. shows a scatterplot of the scores on the first principal component (PC1) plotted against the scores on the second principal comment (PC2). Panel C. shows a scatterplot for the scores on PC1 plotted against the scores on the third principal comment (PC3). Panel D shows a scatterplot of the scores on PC2 plotted against the scores on PC23.

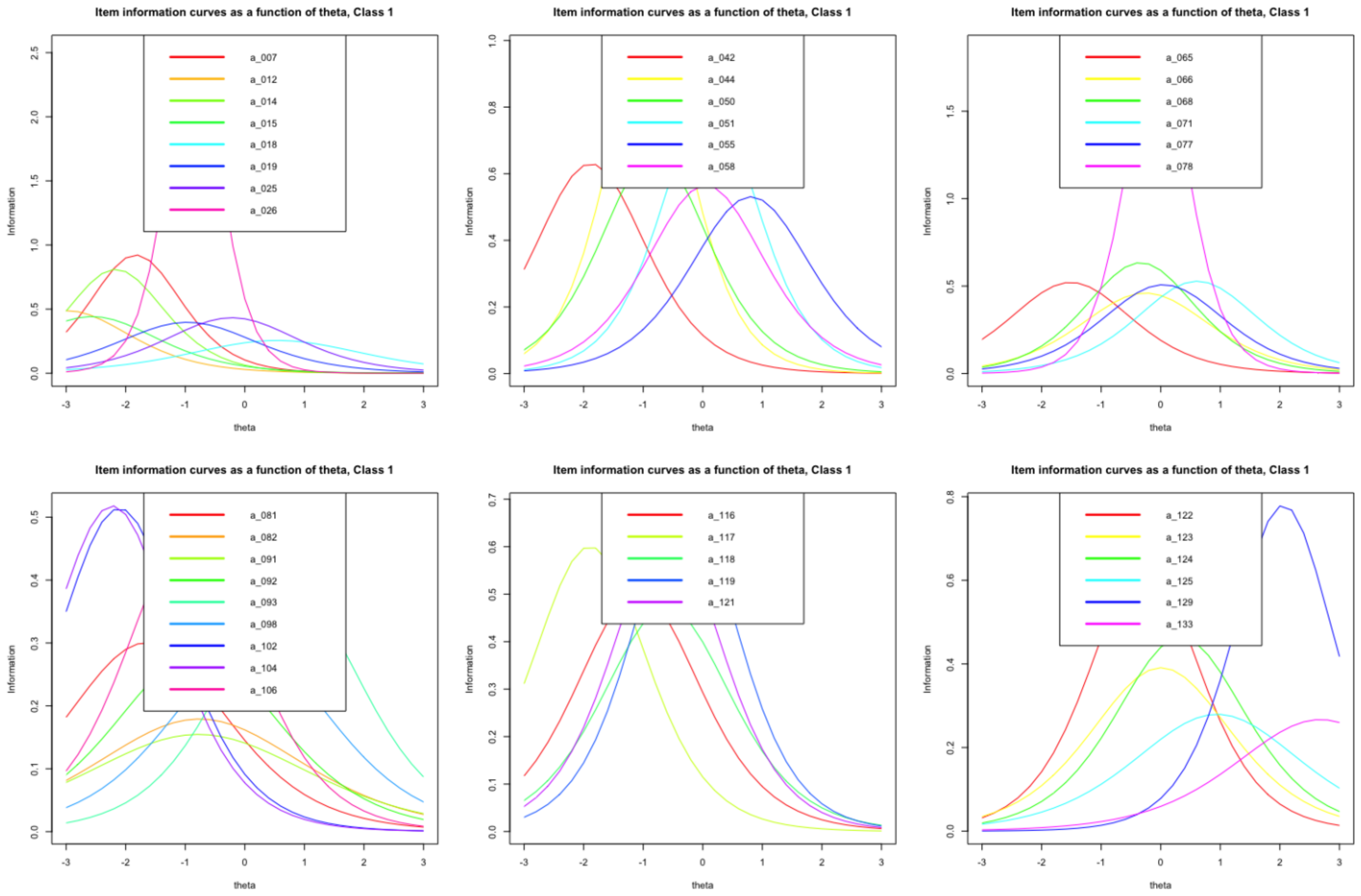


**Supplementary Figure 2**. Item Response Theory (IRT) information curves for the 40 items selected. The top three panels show information curves for the 20 verbal items while the bottom three panels show information curves for the 20 nonverbal items.

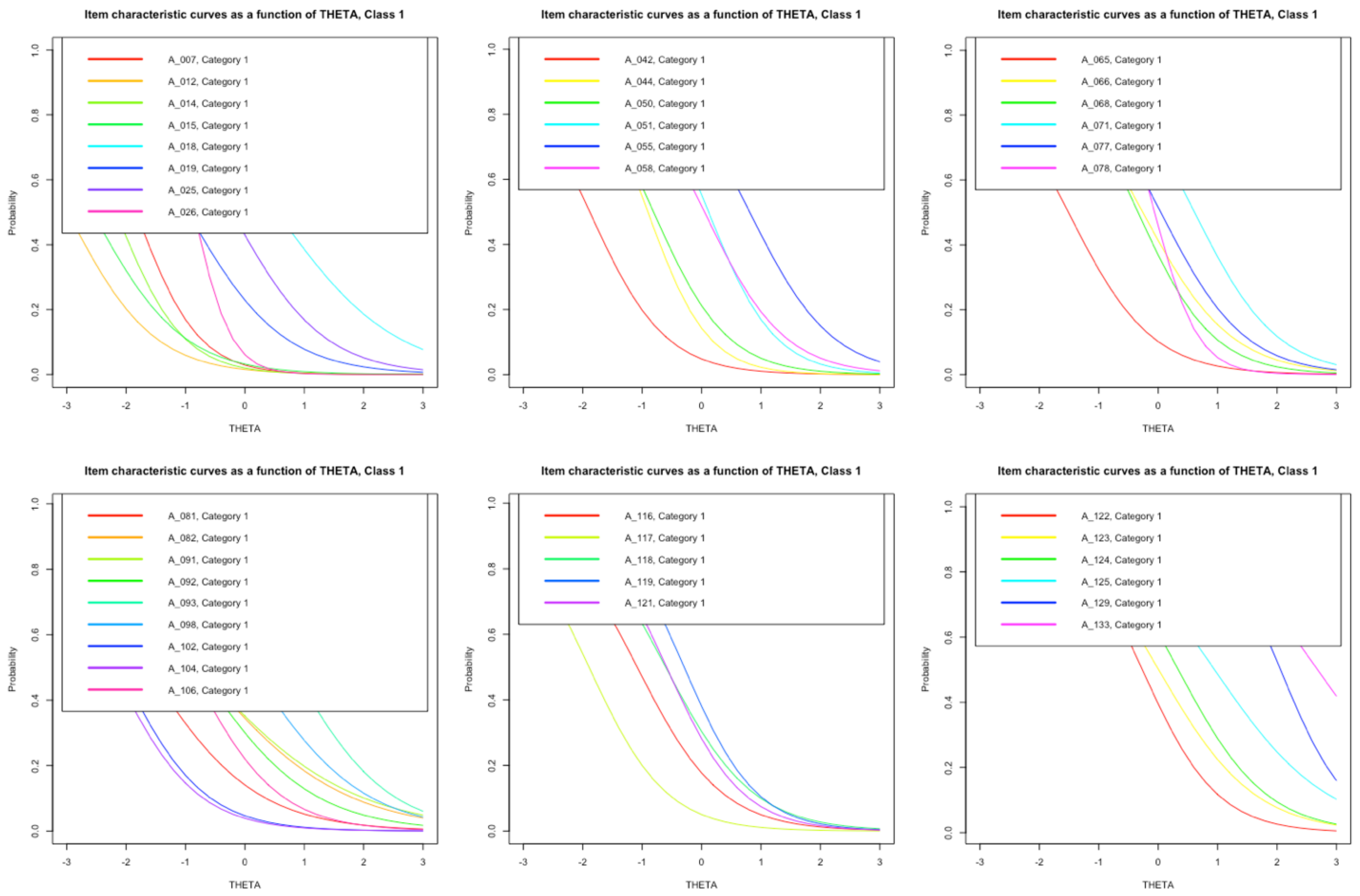


**Supplementary Figure 3**. Item Response Theory (IRT) characteristic curves for the 40 items selected. The top three panels show characteristic curves for the 20 verbal items while the bottom three panels show characteristic curves for the 20 nonverbal items.

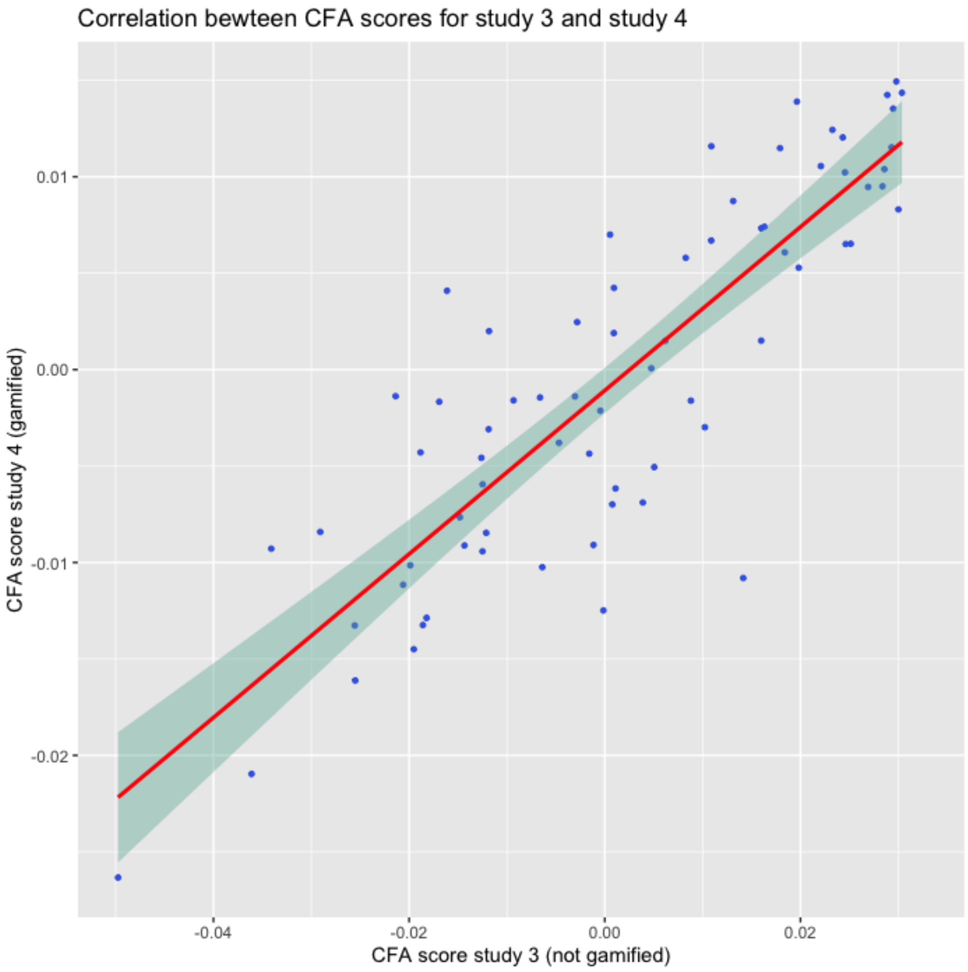


**Supplementary Figure 4.** Correlation (r = .86, p< .0001) between the common factor scores for study 3 and study 4 obtained by including every item collected in study 3 (N = 40, not gamified) and study 4 (N=40, gamified) in two one-factor confirmatory factor analysis (CFA).

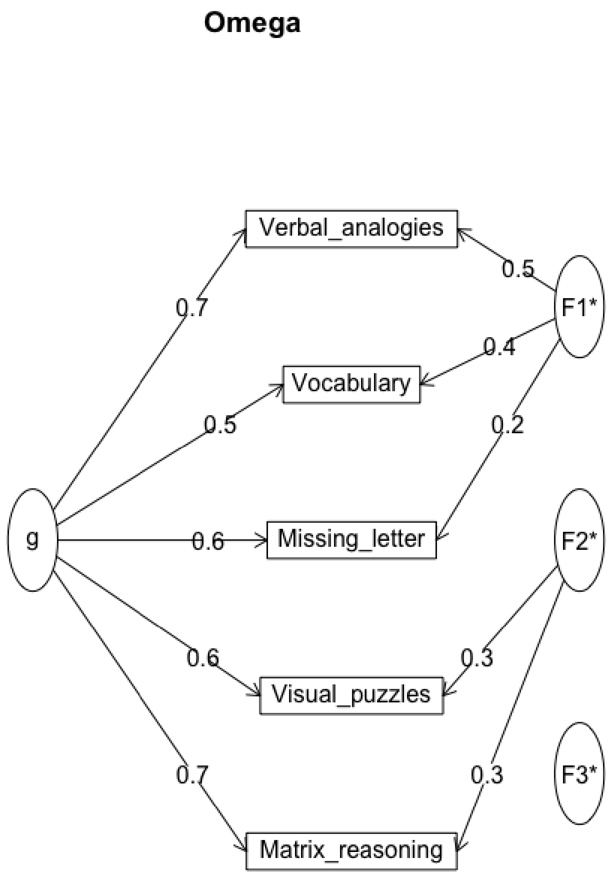


**Supplementary Figure 5.** Path diagram for the g factor and two residual verbal and nonverbal factors obtained using the omega function part of the R package psych (Revelle, 2021).
